# Automated High-Content, High-Throughput Spatial Analysis Pipeline for Drug Screening in 3D Tumor Spheroid Inverted Colloidal Crystal Arrays

**DOI:** 10.1101/2025.06.10.658844

**Authors:** Hyunsu Jeon, Gaeun Kim, James Carpenter, Yamil Colón, Yichun Wang

## Abstract

High-content, high-throughput (HCHT) screening platforms are essential for drug discovery, yet conventional 2D assays lack physiological relevance, and current 3D spheroid systems often face challenges to scalability, uniformity, and the analytical efficiency required for statistically robust screening. Here, we present a fully integrated 3D HCHT platform that synergizes tumor spheroid arrays generated from a bioinert inverted colloidal crystal (iCC) hydrogel framework with an automated, high-speed image analysis pipeline for rapid and spatially resolved therapeutic profiling. The iCC framework enables spontaneous self-assembly of highly-ordered tumor spheroid array at high spheroid density (∼79.8 spheroids·mm^*−*2^) with tight size uniformity (<10% standard deviation), supporting reproducible and high-content imaging (∼40 spheroids·image^*−*1^). The automated image-processing algorithm achieves robust region-of-interest segmentation, fluorescence-weighted centroiding, and multi-parametric spatial analysis in <5 seconds per image—markedly faster than conventional well-plate (5 min per 96 throughput) or single-spheroid analysis workflows methods (∼1 sec per spheroid). Using this platform, we capture dose-dependent diffusion of doxorubicin, tumor penetration profile of small extracellular vesicles, and cell-type-specific infiltration behaviors of monocytes *versus* macrophages. Comparative viability profiling across chemotherapeutics further reveals distinct spatial toxicity signatures, highlighting the importance of spatial context in drug response assessment. Collectively, this platform enables rapid, reproducible, and spatially informative screening in 3D, offering a powerful tool for drug discovery, tumor modeling, and immunotherapeutic development.

## 1. Introduction

Despite an annual global investment exceeding $276 billion—including over $73 billion from companies in the drug development and drug screening,^1^ the field continues to grapple with the challenge of translating preclinical findings into clinical success. High-content, high-throughput (HCHT) drug screening has traditionally relied on twodimensional (2D) culture models^2,3^ due to their operational simplicity, assay compatibility, and ease of scale-up for molecular tracking,^4^ cytotoxicity testing,^5^ and omics-based screening.^6^ Yet, despite their utility, 2D systems fall short in recapitulating key features of native tissue architecture, such as cell–cell and cell–extracellular matrix (ECM) interactions, biochemical gradients, and mechanical signaling, undermining their physiological relevance and predictive accuracy.^7–11^ This has been identified as a major contributor to the persistent bottleneck in translational research: the disconnect between robust preclinical results and successful clinical outcomes.^2,12^ Three-dimensional (3D) culture systems—particularly cellular spheroids, have gained significant momentum as more physiologically representative models. These spheroids self-assemble into compact cell aggregates surrounded by endogenously secreted ECM,^10^ spontaneously forming tissue-like barriers to molecular and mass transport.^13,14^ These spheroid culture models allow spatial signal profiling by enabling quantification in spherical coordinates.^15,16^ This geometrical advantage facilitates 3D *in-vitro* profiling of drug penetration,^17^ nanoparticle trafficking,^18,19^ and immune cell infiltration^20,21^ within the tumor-like microenvironment, representing a critical opportunity to bridge the translational gap between preclinical findings and clinical validations.

However, utilizing spheroid-based models in quantitative HCHT platforms remains technically challenging. For drug discovery and drug screening, an ideal 3D HCHT platform must meet several key criteria: (1) robust and scalable generation of high-yield, uniform spheroids; (2) rapid, efficient, and high-content data acquisition pipelines; and (3) automation-compatible signal analysis and spatial data processing workflows. Although the throughput of spheroid-based screening has improved significantly in recent years,^2,22^ several challenges remain. Traditional techniques such as hanging drop^23^ and spin culture^24,25^ still suffer from variability in spheroid uniformity and yield.^13^ In addition, these methods offer limited compatibility with automated imaging and analysis pipelines,^26,27^ which further hinder the use of spheroid-based models for statistically powered drug screening.^2,28^ To address these limitations, various engineering strategies have been developed. For instance, round-bottom well plates have enabled scaled production in 96-, 384-, and 1536-well formats (yielding one spheroid per well) over the past decade,^29–31^ however, for efficient data acquisition they exhibit spheroid density at only <0.15 spheroids/mm^2^ in a 127.76 mm × 85.48 mm plate. On the other hand, to further improve efficiency of data acquisition, especially for HCHT image-based analyses, custom-fabricated micropillar arrays have been developed to achieve denser spheroid generation (*e*.*g*., 600 spheroids in a 3.3 mm × 11.3 mm area, ∼16.09 spheroids/mm^2^).^32,33^ More recently, the use of biomaterials such as hydrogels has enabled the fabrication of spatially organized spheroid arrays via bioprinting, where their tunable mechanical properties and biocompatibility facilitate precise, simultaneous deposition of multiple spheroids through digitally controlled nozzle arrays (*e*.*g*., ∼6 spheroids in 1 mm^3^).^34^ Despite recent advances, many engineered platforms rely on complex material formulations—such as the incorporation of additional reagents for spheroid formation—that hinder scalability and are not readily compatible with end-to-end 3D HCHT screening workflows.^35^ These material and system-level limitations restrict the deployment of dense spheroid arrays for spatiotemporal high-resolution imaging and robust quantitative signal extraction.

While these advances have improved spheroid yield and compatibility with conventional analysis tools, there is still critical need for downstream automation of spatial signal analysis, a major remaining barrier to full HCHT implementation.^36^ Indeed, although spheroids have been widely used for tracking biological signals spatially (*e*.*g*., marker expression, therapeutic carrier transport, and immune cell invasion),^13,37,38^ most current spatial analysis pipelines still rely heavily on manual or semi-automated image processing (*e*.*g*., manual segmentation and parameter adjustment) for spheroid identification and signal localization.^39–41^ This reliance not only introduces time-consuming workflows and user-dependent variability, but also significantly limits scalability, not only when applying ImageJ plugin (*e*.*g*., Radial Profiler),^42^ but even when supported by commercial or open-source tools like TASI,^39^ Spheroid Analyzer,^43^ or SpheroidSizer.^44^ Furthermore, these tools are typically designed to process images containing a single spheroid, making both data input and output extraction labor-intensive and impractical for large-scale image datasets,^45^ which severely constrain their utility in statistically powered drug screening. To enable HCHT applications with high-yield cell spheroids for broader users, here is a pressing need for robust, automation-compatible analysis pipelines. These pipelines must be capable of processing 3D spatial signals across hundreds to thousands of spheroids per dataset with minimal user intervention. More importantly, they should integrate fully automated segmentation, spatial registration, and quantification workflows.

To overcome these challenges, we developed a bioinert, optically transparent hydrogel-based framework with a tunable physical architecture that enables reproducible, high-density spheroid formation and downstream imaging. This inverted colloidal crystal (iCC) framework leverages electrosprayed alginate microgels as a sacrificial template^46^ to form a highly ordered porous architecture within a non-adhesive, mechanically robust, and biocompatible agarose matrix, ^47^ which facilitates spontaneous and uniform spheroid self-assembly organized in a crystalline array.^48^ This framework enables the spontaneous formation and crystalline arrangement of uniform tumor spheroids in high yield (∼79.8 spheroids/mm^2^). Building on this biomaterial-engineered iCC platform, we developed a fully automated, material-integrated methodology for HCHT spatial analysis, enabling advanced applications in 3D spheroid-based drug screening. This scalable, optically transparent system provides a robust and reproducible 3D platform, tumor spheroid iCC arrays, for spatially resolved assessment of chemotherapeutic transport, nanocarrier delivery, immune cell infiltration, and treatment-induced viability changes. Importantly, the spatioperiodic architecture of tumor spheroid iCC array enhances experimental reproducibility, both radially within individual spheroids and across biological replicates, enabling consistent HCHT analysis. To realize front-to-end automation in 3D HCHT pipeline, we developed a custom algorithm, capable of region of interest (ROI) recognition, coordinate-based spheroid extraction, fluorescence-weighted centroid detection, and vectorized radial signal profiling. The pipeline can process each image—containing ∼40 spheroids—in under 5 sec, significantly enhanced the processing time compared to conventional methods, such as well-plate-based automated analysis (5 min per 96 throughput)^49^ or individual spheroid profiling (∼1 sec per spheroid).^50^ The integration of our imaging and analysis pipeline with 3D tumor spheroid arrays embedded in the iCC framework enables rapid, quantitative spatial profiling of drug penetration, nanoparticle trafficking, and immune cell (monocyte/macrophage) migration within the HCHT 3D environment. Utilizing this system, we reveal unconventional therapeutic penetration behaviors, indicating the challenges in nanoparticle diffusion and the role of active immune cell transport. Additionally, the platform enables spatial toxicity mapping through adaptive image thresholding. Chemotherapy profiling of cisplatin, doxorubicin, and paclitaxel in this system uncovered distinct, drug-specific viability gradients, illustrating the differential behavior of small-molecule drugs within 3D tumor tissue. Together, this integrated, automation-ready system offers precise and scalable capabilities for advancing 3D HCHT applications in cancer biology, drug discovery, and therapeutic evaluation.

## 2. Result

### 2.1 Tumor Spheroid Array Generation

To enable HCHT generation of spatially organized 3D tumor spheroids as an array, we constructed a bioinert iCC hydrogel framework featuring a mechanically robust, hexagonally packed architecture, initially developed in our work.^48^ The framework features a crystalline structure composed of uniformly distributed spherical cavities (*e*.*g*., termed void spaces; ∼250 µm in diameter) interconnected by narrow channels (*e*.*g*., ∼50 µm) (**Figure 1a**), serving as a mechanically robust and structurally organized platform for generating highly ordered and size-controlled spheroid arrays. This framework was created by a scalable template, hexagonally close-packed (HCP) alginate microgels with colloidal crystal (CC) structure assembled *via* a pseudo-isolated slit dehydration method (*i*.*e*., sonication-assisted dehydration in a water-selective pump-out slit; **Figure 1b**), which also enable gravity-mediated overlapping of microgels due to their viscoelastic properties. The microgel template was embedded in a thermos-reversible agarose hydrogel^51,52^ and then removed through enzymatic degradation and calcium chelation.^46,48^ The resulting iCC framework contained openings to upper void spaces, enabling uniform cell seeding *via* perpendicular centrifugation, while its low-adhesive environment provided precise spatial confinement conductive to individual spheroid generation (**Figure 1c**). Representatively, an iCC tumor spheroid array composed of size-controlled hepatocarcinoma (HepG2) spheroids formed spontaneously upon centrifugal cell seeding high yield and uniformity (*e*.*g*., >50 spheroids·μL^−1^; 161.1 ± 12.8 µm; **Figure S1**) while maintaining HCP arrangement of them (*e*.*g*., See left panel; **Figure 1e**). This setup enabled scalable, on-demand two-photon confocal laser scanning microscopy (TP-CLSM), ranging from HCHT imaging of hundreds of spheroids per scan to single-spheroid resolution (**Figure 1e**). Thus, our setup provided consistent optical accessibility with high imaging throughputs (*e*.*g*., >50 spheroids·image^−1^ in < 4 min·image^−1^) for HCHT data acquisition, collection, and spatial analysis. Altogether, highly uniform, low-variability spheroid arrays were established through the iCC framework, providing an ideal and robust foundation for automated, multiplexed analysis in HCHT drug screening and mechanistic studies.

**Figure 1.**
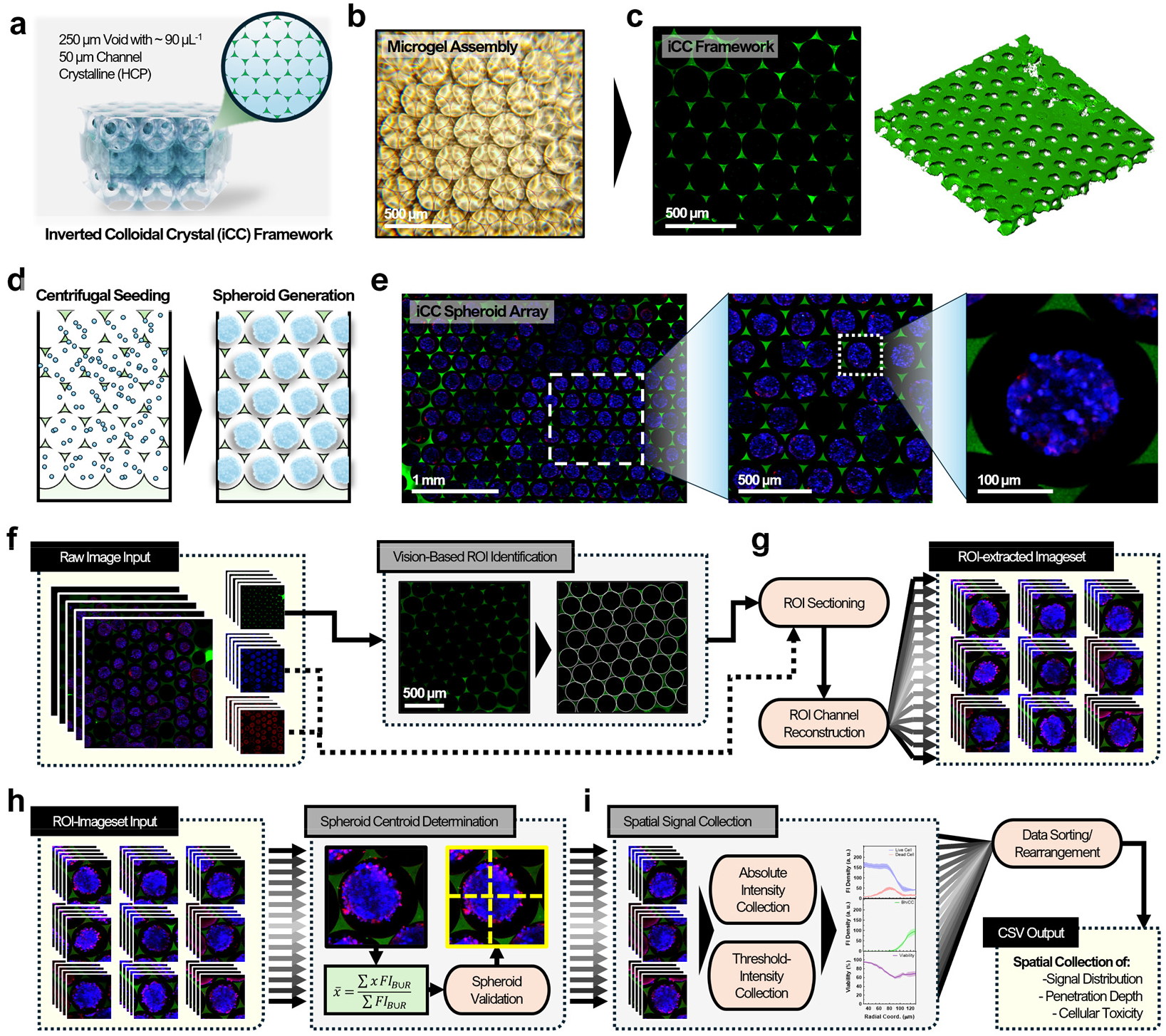
Inverted colloidal crystal (iCC) spheroid array generation, imaging, and automated data processing workflow. **(a)** Schematic of the bioinert iCC framework with 250 µm voids and 50 µm interconnected channels for high-yield spontaneous spheroid formation. **(b)** A bright-field microscopic image of alginate microgel assembly. **(c)** A representative two-photon confocal laser scanning microscopy (TP-CLSM) image of the iCC framework. (**d**) Schematic of the spheroid array generation *via* centrifugal cell seeding into the iCC framework. (**e**) Representative TP-CLSM images of spheroid arrays in crystalline arrangement. **(f)** The automated region of interest (ROI) identification and **(g)** extraction from raw multichannel TP-CLSM images. **(h)** Spheroid centroid determination with fluorescence-weighted mean and **(i)** spheroid-level signal quantification with spatial profiling for intensity-based analyses of distribution, penetration depth, and cellular toxicity.

### 2.2 Automated Data Processing Algorithm Development

To effectively and efficiently process the HCHT TP-CLSM imaging data from the iCC spheroid array—containing ∼40 spheroids per image in this study, with scalable capacity in imaging—an automated data processing algorithm was developed. This algorithm enables individual spheroid recognition, spherical coordinate system assignment, and spatial data collection. The overall workflow consists of two main parts: (i) recognizing and segmenting the raw multichannel images into individual spheroid-specific regions across all channels (**Figure 1f-g**), and (ii) determining and analyzing each spheroid in its own coordinate system (*i*.*e*., spherical coordinates for each spheroid; **Figure 1h-i**).

To identify and extract void spaces in iCC framework as primary regions of interest (ROIs) from raw multichannel TP-CLSM images—comprising green fluorescence from the framework and blue/red fluorescence for spheroid and target signal tracking—we implemented an opensource circle detection algorithm on the green channel (*e*.*g*., Hough Circle Detection^53^, OpenCV; **Figure 1f**). Applied to the iCC framework signal, this algorithm accurately located the Cartesian coordinates of individual circular voids (**Figure S2**), enabling automated segmentation and storage of ROI coordinates for downstream processing (**Figure 1g**).

Fluorescence signals within each segmented ROI were further analyzed using fluorescence-weighted centroiding (**Figure 1h**). The centroid coordinate was calculated the central point of each spheroid based on cellular signals, thus establishing the spherical coordinate origin for each spheroid (**Equations (1)–(2)**). This centroiding approach minimizes discrepancies between the ROI origin and the actual spheroid center, which is particularly advantageous for spatially resolved signal profiling in 3D spheroids, as it enables precise localization of signal gradients relative to the spheroid center. Finally, by applying linear algebra-based vectorization, spatial signal extraction in spherical coordinates (**Equation (3)**) was achieved with significantly improved processing efficiency (*e*.*g*., > 17.508 sec per image with ∼40 spheroids; **Figure 1i**), compared to the conventional for-loop-based 3D sorting method for the same images (*e*.*g*., < 30 sec per image). This approach accommodated both raw and thresholded intensity values and was compatible with downstream quantitative modeling, including spatial viability estimation, radial dimension normalization, and data representation.

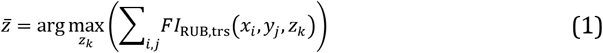

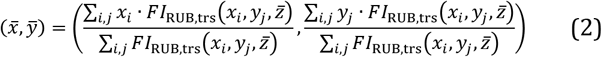

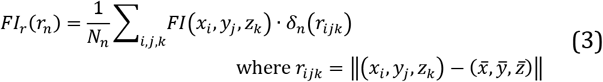

Altogether, this integrated pipeline significantly advances 3D HCHTS enabled by the iCC spheroid array. It offers fully automated ROI detection, centroid determination, and spatial signal profiling. Unlike traditional methods that rely on manual or semi-manual segmentation—often hindered by densely packed spheroids and indistinct ROI boundaries^39–41,54^—this methodology delivered rapid processing speeds (*e*.*g*., <10 sec·image^−1^ with personal laptop (i7-1165G7; 16 GiB RAM; 1TB NVMe SSD; Iris Xe Graphics) and <5 sec·image^−1^ with Computational Facility (Xeon(R) Gold 6326; 256 GiB; Networked AFS; 4 x NVIDIA A10); **Table S1**), supporting the automated 3D HCHT operation of our system. On average, each image contains ∼40 spheroids, and with scalable imaging capacity up to ∼3000 spheroids per sample group (*e*.*g*., increased scan size in TP-CLSM), enabling streamlined processing of thousands of spheroids across hundreds of images with minimal user input. Additionally, the pipeline incorporated validation steps, including spheroid existence checks and size/location-based qualification, ensuring robustness and accuracy across large datasets.

### 2.3 3D HCHT Spatial Profiling of Doxorubicin in HepG2 Spheroid Array

To evaluate and validate our spatial signal processing algorithm, we first assessed the penetration of doxorubicin within the iCC-assembled HepG2 spheroid array using TP-CLSM (**Figure 2a**). For accurate spatial profiling, the full extent of each spheroid was defined by OR-gating the blue (*i*.*e*., live cells) and red (*i*.*e*., dead cells) fluorescence channels, ensuring complete capture of both viable and non-viable HepG2 regions. Doxorubicin distribution was detected in the green channel,^55^ enabling spatial colocalization analysis relative to the spheroid structure. **Figure 2b** presents representative multichannel TP-CLSM images of HepG2 spheroids (magenta) exposed to increasing concentrations of doxorubicin (green). At lower doses (10 μM), doxorubicin fluorescence appeared faint and was primarily localized to the spheroid periphery, with minimal overlap with the cellular signal in the central regions — suggesting limited penetration at a low dose. As the concentration increased to 50 and 200 μM, the drug signal became more homogeneously distributed throughout the spheroid volume, exhibiting greater colocalization with the spheroid. This dose-dependent spatial shift highlighted the progressive penetration of doxorubicin into the spheroid interior. To quantitatively evaluate this trend, we employed our automated data processing algorithm with raw intensity collection under absolute radial dimension to the TP-CLSM images; **Figure 2c** displays the corresponding radial intensity profiles of both doxorubicin and cell signals across spheroid cross-sections. These plots confirmed that lower doxorubicin concentrations exhibited steep signal gradients with peripheral localization, whereas higher concentrations resulted in flatter radial distributions, indicative of deeper and more uniform penetration. Together, these quantitative profiles validated the qualitative observations in **Figure 2b** and demonstrated the utility of the iCC framework-based 3D HCHTS platform for high-information-content, spatially resolved drug assessment across multiple spheroids in parallel (*i*.*e*., ∼40 spheroid per image).

**Figure 2.**
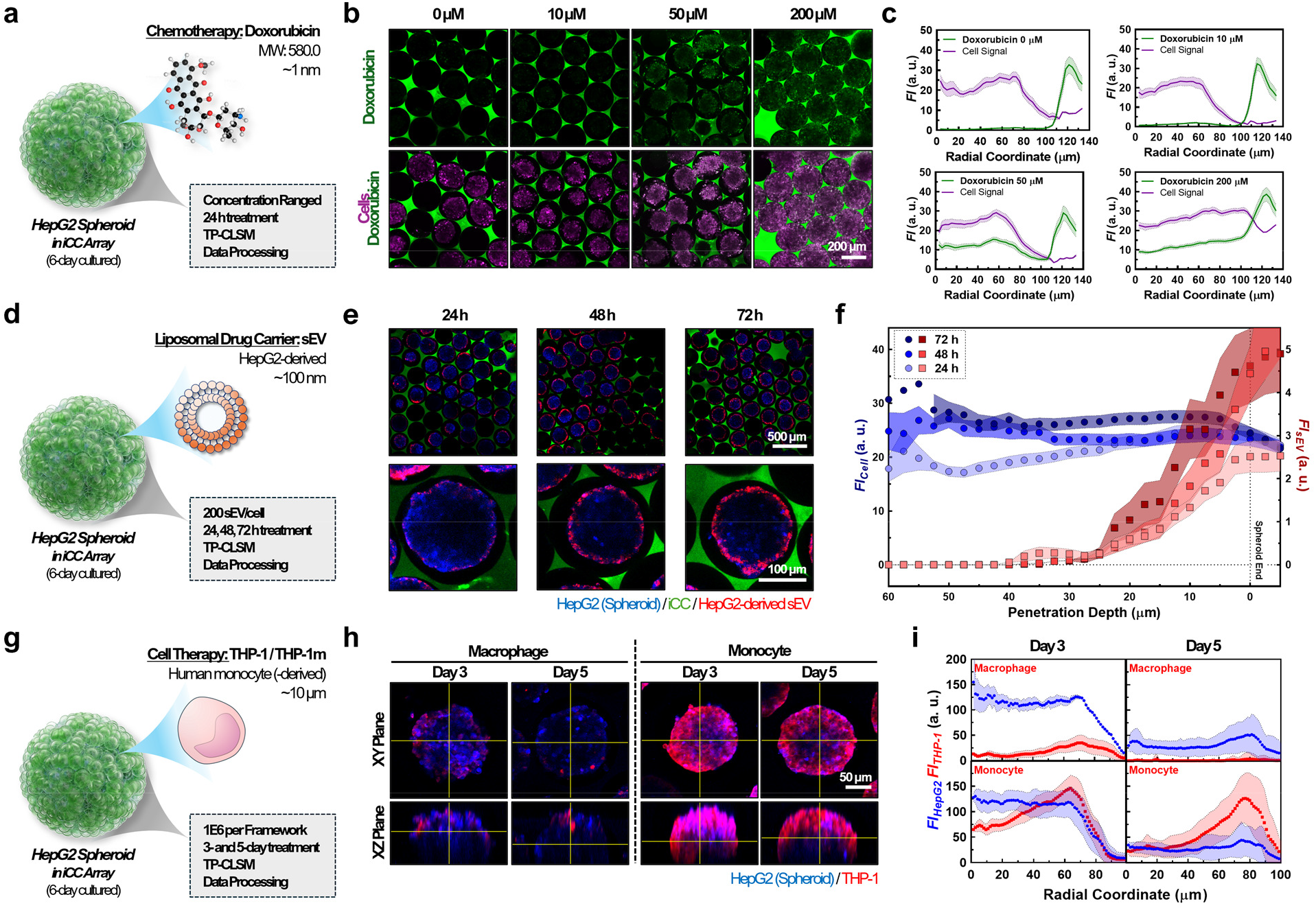
Spatial signal profiling of **(a-c)** doxorubicin, **(d-f)** small extracellular vesicles (sEVs), and **(g-i)** THP-1 monocyte/macrophage (THP-1/THP-1m) penetration within the iCC spheroid array platform. **(a)** Schematic of doxorubicin tracking test in the iCC spheroid array. **(b)** Representative TP-CLSM images showing dose-dependent doxorubicin (green) penetration in HepG2 spheroids (magenta: merged Blue Cell Tracker and BOBO-3) across increasing concentrations (0–200 µM). **(c)** Corresponding radial signal profiles of doxorubicin and cell intensity across spheroid crosssections at each dose. **(d)** Schematic of sEV tracking test in the iCC spheroid array. **(e)** Time-lapse imaging of HepG2derived sEVs (red) trafficking into HepG2 spheroids over 24, 48, and 72 hours. **(f)** Quantification of average cell signal (blue, left y-axis) and sEV signal (red, right y-axis) across spheroid radial coordinates over time. **(g)** Schematic of THP-1/THP-1m tracking test in the iCC spheroid array. **(h)** Orthogonal (XY/XZ) projections of HepG2 spheroids (blue) coincubated with THP-1–derived macrophages (left) or monocytes (right), showing differential penetration of THP-1m or THP-1 (red) at Day 3 and Day 5. **(i)** Spatial fluorescence profiles comparing THP-1m/THP-1 and HepG2 spheroid signal distributions at Day 3 and Day 5.

### 2.4 Automated Data Processing Algorithm Development

To expand the applicability of the iCC spheroid array and its data processing algorithm in 3D HCHTS, we broadened the scope beyond small chemical drugs (*e*.*g*., doxorubicin) by including lipid-based drug carrier (small extracellular vesicles, sEVs)^56^ and immune cells (THP-1 monocyte/macrophage; THP-1/THP-1m)^57,58^ that are increasingly recognized as promising candidates in drug delivery and cell therapy.^59,60^ We utilized the iCC spheroid array system and the automated data processing algorithm to gather sEV penetration and transport profiles over time, establishing their spatiotemporal dynamics. The sEVs were isolated from conventional HepG2 cell cultures (Hep-sEVs) using our established protocol.^61,62^ Transmission electron microscopy (TEM) confirmed the spherical morphology and structural integrity of the isolated Hep-sEVs (**Figure S3a**). Nanoparticle tracking analysis (NTA) revealed a concentration of (6.62 ± 0.56) × 10^?^ particles/mL and a narrow size distribution of 114.8 ± 5.8 nm (**Figure S3b**). The narrow size distribution corresponded to a negative surface charge of −8.21 ± 1.16 mV (**Figure S3c**), suggesting excellent colloidal stability of the isolated sEVs. To further validate the identity of the isolated nanoparticles as sEVs, the expression of key tetraspanin biomarkers (CD9, CD63, and CD81) was confirmed through Western blot analysis (**Figure S3d**).

To investigate the spatiotemporal dynamics of Hep-sEV penetration (*e*.*g*., ∼100 nm size range) within the HepG2 spheroid array, we treated 6-day cultured iCC-assembled HepG2 spheroids with DiD-stained Hep-sEVs at 24, 48, and 72-h time points (**Figure 2d**). **Figure 2e** presents representative TP-CLSM images of the spheroid arrays and zoomed in region for individual spheroid at each time point, with the blue channel for live cells and the red channel for sEVs. A limited penetration of sEV (*e*.*g*., ∼20 µm) was observed, with the red signal accumulating primarily at the spheroid edges. No significant progression of sEV penetration into the spheroid interior was observed, even after 72 h of treatment, suggesting that the sEVs were unable to penetrate deeper into the spheroid across the entire array. The data processing algorithm provided a more quantitative representation, as shown in **Figure 2f**. The processed spatial (radial coordinate) curves of fluorescence intensities from live cell (*Ff*_*Cell*_) and sEV (*Ff*_*sEV*_) quantified the penetration of sEVs within tumor spheroids over time. While the *Ff*_*Cell*_ curves (*e*.*g*., blue theme) showed a plateau across all penetration depths and treatment times, the *Ff*_*sEV*_ curves (*e*.*g*., red theme) revealed relatively high intensity values accumulated at the spheroid edges (*i*.*e*., 0-20 µm penetration depth), with minimal to negligible intensity observed in the interior at all time points. These time-lapse qualitative and quantitative 3D HCHT results from iCC spheroid array emphasize the penetration barrier of Hep-sEVs in 3D tumor environments generated by HepG2 liver cancer cells.

To further validate the iCC spheroid array and its automated analysis system, we investigated the spatiotemporal dynamics of immune cell penetration within the spheroid array, using macrophages (THP-1m) and monocytes (THP-1) over a 3, 5, and 7-day treatment period (**Figure 2g**). The THP-1 cell line, a human monocytic model, is widely used to study monocyte migration and activation in tumor tissues,^59^ while THP-1m macrophages are commonly used to simulate immune cell behavior in the tumor microenvironment, where they either exert both tumor-suppressive or tumor-promoting effects.^63^ THP-1 cells were differentiated into THP-1m macrophages *via* PMA treatment (*e*.*g*., 100 ng·mL^-1^ for 24 h, followed by 24 h of incubation in fresh media) and subsequently stained with a red cell tracker (*i*.*e*., CMTPX). **Figure 2h** shows representative TP-CLSM images in XY planes (single slice) of the iCC spheroid array treated with THP-1m or THP-1 cells (top) and its orthogonal projections in XZ planes (bottom). Notably, the immune cell signal (red channel) was significantly more pronounced in THP-1 cells than in THP-1m, highlighting cell-specific differences in penetration behavior. Quantitative analysis using our automated algorithm (**Figure 2i)** revealed distinct differences in the penetration profiles and localization of the two cell types. THP-1m macrophages displayed slight infiltration on Day 3, with most of the signal from THP-1m disappearing by Day 5, likely due to cytokine-induced cell death.^64,65^ In contrast, THP-1 monocytes showed more substantial and consistent signal distribution throughout the spheroid, with a gradual increase in fluorescence intensity at the spheroid edges on both Day 3 and Day 5. These results emphasize the distinct penetration behaviors between two types of immune cells within a 3D tumor-like microenvironment. Importantly, this platform and corresponding analysis algorithm demonstrated their capacity to distinguish immune cell dynamics in 3D HCHTS, indicating their great potential for studying immune cell behavior and cell therapies within 3D tumor cultures with high statistical reliability.

### 2.5 3D HCHT Spatial Profiling of Small Chemical Drug-Mediated Cellular Viability in the iCC Spheroid Array

To further demonstrate the versatility of the iCC spheroid array system and its automated analysis pipeline for drug screening applications, we adapted our algorithm to enable spatially resolved quantification of cellular viability. As proof-of-concept, three commonly used small-molecule chemotherapeutics—(1) doxorubicin, (2) cisplatin, and (3) paclitaxel—were selected for treatment in the iCC spheroid array platform. Prior to spheroid generation, HepG2 cells were pre-stained with a blue live-cell tracker that is cell-permeant, retained in the cytoplasm, and fluoresces upon reaction with intracellular primary amines without binding to specific organelles or structures. The spheroids were then cultured for 6 days, followed by a 24-h drug treatment. After treatment, spheroids were stained with a red dead-cell indicator (*e*.*g*., BOBO-3; a membrane impermeant nucleic acid stain), fixed, and imaged using TP-CLSM for raw data acquisition, followed by our spatial analysis algorithm processing. Importantly, spatial viability was calculated based on thresholded fluorescence intensities of live and dead cell channels (*e*.*g*., *Ff*_*B,trs*_ and *Ff*_*R,trs*_ ; see **Equation (4)** and *Experimental Section*), allowing for precise quantification of dead cell distribution across the full spheroid volume on a ROI-specific basis.

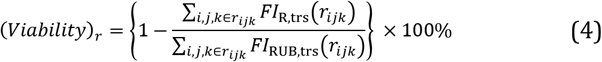

First, we conducted a dose-dependent HCHT spatial profiling study of HepG2 spheroids treated with doxorubicin. Doxorubicin, an anthracycline-based chemotherapeutic with a molecular size of approximately 1 nm and a molecular weight of ∼580 Da, is widely known for its DNA intercalation and reactive oxygen species (ROS)-mediated cytotoxicity.^66,67^ In conventional 2D cultures, doxorubicin exhibits an *fC50* of ∼1.3 µM (denoted as *fC50*_*2D*_)^68^ for HepG2 cells after 24 h of treatment (**Figure 3a**). Following 24 h treatment with doxorubicin at concentrations ranging from 5 to 500 µM, *Z*-projected TP-CLSM imaging (**Figure 3b**) qualitatively depicted live (blue) and dead (red) cell populations. At concentrations starting from 20 µM, a notable drop in cell viability was observed, with the red signal becoming dominant across the spheroids in the iCC spheroid arrays, particularly at higher doses (50–500 µM). To gain deeper insights into the spatial patterns of viability loss quantitatively, our automated algorithm processed the raw dataset, generating both representative frequency heatmaps (**Figure 3c**) and individual plots (**Figure S4**) of spatial cell viability. These heatmaps and plots were mapped across radial coordinates and viability percentiles, providing a spatial and quantitative analysis of cellular response to doxorubicin treatment. While low doses (*e*.*g*., 5 µM) induced subtle, radially shallow reductions in viability, higher doses (*e*.*g*., ≥50 µM) resulted in near-complete, plateaued cell death across the spheroid volume. Polar plots of spatial viability (**Figure 3d** and **Figure S5a**) further illustrate the spatial discrepancy between the spheroid core (*i*.*e*., denoted as *0*.*0R*) and edge (*i*.*e*., denoted as *1*.*0R*) regarding dose- and depth-dependent reductions in cell viability. Low doses presented distinct differences in spatial viability between the core and edge (*i*.*e*., *eViability* _*1*.*0R*_ *− eViability* _*0*.*0R*_ at 5 and 20 µM across polar coordinates; **Figure S5b**), while the difference gradually diminished as drug concentration increased. Meanwhile, the cell viability in higher doses (*e*.*g*., ≥50 µM) was nearly all-zeroed across the radius of spheroids, aligning with both heatmap and image-based observations in **Figure 3b–c**.

**Figure 3.**
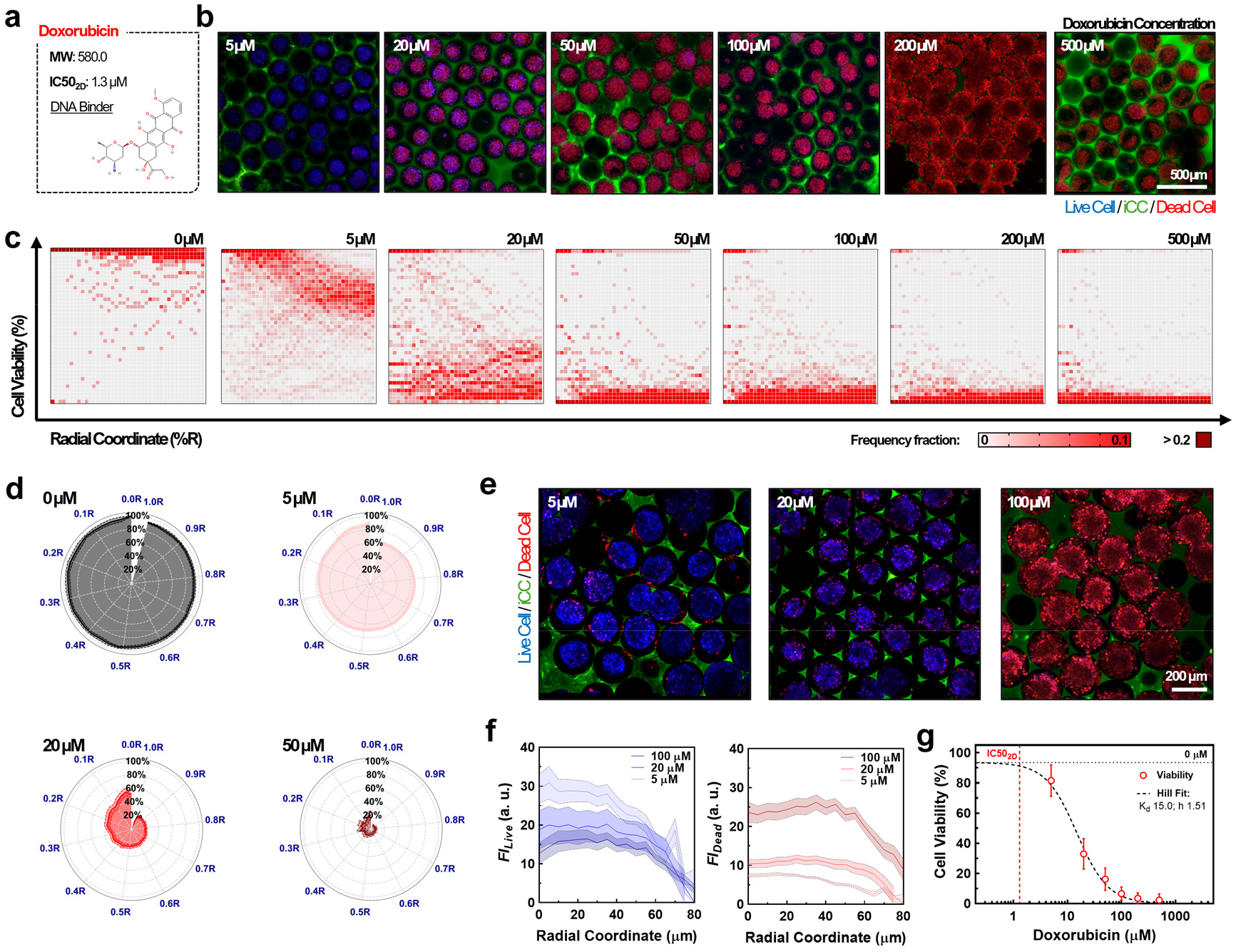
HCHT spatial viability profiling of HepG2 spheroids exposed to doxorubicin in the iCC spheroid array (24 h treatment following 6-day spheroid culture). **(a)** Descriptive summary of doxorubicin. **(b)** Representative TP-CLSM images showing live (blue) and dead (red) HepG2 cell distributions in response to increasing doxorubicin concentrations (5–500 µM). Green: iCC framework. Images represent sum slice-based Z projections. **(c)** Heatmaps of cell viability frequency across radial coordinate (%R) and viability percentile at varying doses. **(d)** Polar plots visualizing spatial viability distributions at 0, 5, 20, and 50 µM, highlighting dose- and depth-dependent reductions in live cell populations. **(e)** Single-slice TP-CLSM images showing radial viability changes at 5, 20, and 100 µM. **(f)** Radial fluorescence intensity profiles for live (left) and dead (right) cell signals, showing a plateaued viability drop at selected concentrations. **(g)** Dose–response curve with IC50 estimation *via* Hill fitting.

To further investigate the spatial viability trends from doxorubicin treatment in the iCC spheroid array, we revisited the raw TP-CLSM images shown in **Figure 3e**, which present single-*Z*-slice images at representative doses (5, 20, and 100 µM) with more straightforward representation of *Ff* distribution. Consistent with the results from **Figure 3b-d**, at lower doses, dead cells were primarily localized at the spheroid edges, contributing to the observed radial viability patterns. However, at higher concentrations, distinct edge-localized patterns were lost, with cell death occurring more homogenously throughout the spheroid. *Ff* profiles of live and dead signals (**Figure 3f**) demonstrated that *Ff*_*B*_ (*i*.*e*., live cell) decreased progressively with increasing doxorubicin concentrations, while *Ff*_*R*_ (*i*.*e*., dead cell) increased across radial depths. Notably, significant enhancement of dead cell signals at the spheroid edge was primarily observed at the 5 µM dose, with a slight trend at 20 µM. However, at higher concentrations, the signal profiles plateaued, indicating a more uniform distribution of cell death throughout the spheroid. These dose-responsive patterns are spatially comparable to the results observed in **Figure 2a-b** with intensity-based spatial penetration assessments, showing lower dose but display deeper infiltration of effect due to the smaller molecular size of doxorubicin.

To quantify the overall cytotoxic response, total cell viability calculated from **Equation 5** was plotted against drug concentration and fitted using the Hill equation for IC_50_ estimation (**Figure 3g**). The resulting curve and Hill fit showed higher *fC50* from spheroids (denoted as *fC50*_*3D*_) as ∼15 µM than prior literature-reported responses of HepG2 to doxorubicin (*fC50*_*2D*_∼1.3 µM^68^), supporting the certain transport limitation in lower doses of doxorubicin into HepG2 spheroids.

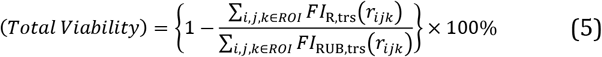

Next, we investigated the spatial cytotoxic response of HepG2 spheroids to cisplatin, a platinum-based DNA crosslinker with a molecular weight of approximately 300 Da (**Figure 4a**). In conventional 2D cultures, cisplatin exhibits an *fC50*_*2D*_ of ∼25.5 µM^69^ for HepG2 cells after 24 h of treatment—approximately 20-fold higher than the *fC50*_*2D*_ of doxorubicin, indicating its lower potency in monolayer conditions. To accommodate this difference and reflect solubility constraints, the cisplatin concentration range for the iCC spheroid array was selected from 50 µM (*e*.*g*., 10-fold higher than the lowest dose used for doxorubicin) to a maximum of 2000 µM, the upper solubility limit. Following 24 h of treatment, sum-sliced TP-CLSM imaging (**Figure 4b**) qualitatively revealed that even at the highest applied concentration, live-cell signals (blue) remained prominent, indicating incomplete cytotoxicity, while dead-cell signals (red) began to accumulate, particularly at the spheroid periphery. This trend was further supported by spatial frequency heatmaps (**Figure 4c**; individual plots in **Figure S6**), which quantitatively showed edge-initiated viability loss that became progressively more pronounced with increasing doses. Notably, a visible reduction in peripheral viability was evident from 200 µM, while toxicity gradually appeared toward the spheroid core at higher concentrations. Polar plots (**Figure 4d**) demonstrated persistent radial discrepancies in viability (**Figure S7a**), with *eViability* _*1*.*0R*_ *− eViability* _*0*.*0R*_ maintained across all doses. The most significant gap (up to ∼50% difference in viability; **Figure S7b**) was observed at 1000 µM with statistical significance, indicating that edge-to-core viability gradients were preserved even under the highest dose we can treated. This spatial trend was consistently observed in TPCLSM images (**Figure 4e**) and further supported by *Ff* profiles across radial coordinates (**Figure 4f**). At lower doses (<100 µM), live cells remained predominantly viable within the spheroid core. However, at higher doses (400 µM and 1000 µM), live-cell signals (*Ff*_*B*_) diminished throughout all region, while dead-cell signals (*Ff*_*R*_) became increasingly concentrated at the periphery, indicating substantial edge-specific cytotoxicity. Notably, dead signal profiles peaked near the spheroid edge at high-dose treatment, contrasting with doxorubicin treatment which showed plateaued profiles (**Figure 3**). To quantify overall cytotoxicity, total cell viability was plotted as a function of drug concentration and fitted using the Hill equation (**Figure 4g**), yielding an estimated IC_50_ value in 3D platform (*fC50*_*3D*_**∼**883 µM). This is markedly higher than reported *fC50*_*2D*_ ∼25.5 µM in traditional 2D platform,^69^ reflecting a ∼40-fold increase. This discrepancy is potentially attributed to the limited drug penetration into the spheroid core during cisplatin treatment, leading to delayed cytotoxic effects; the elevated *fC50*_*3D*_ may reflect reduced intratumoral diffusion and spatially variable drug exposure.

**Figure 4.**
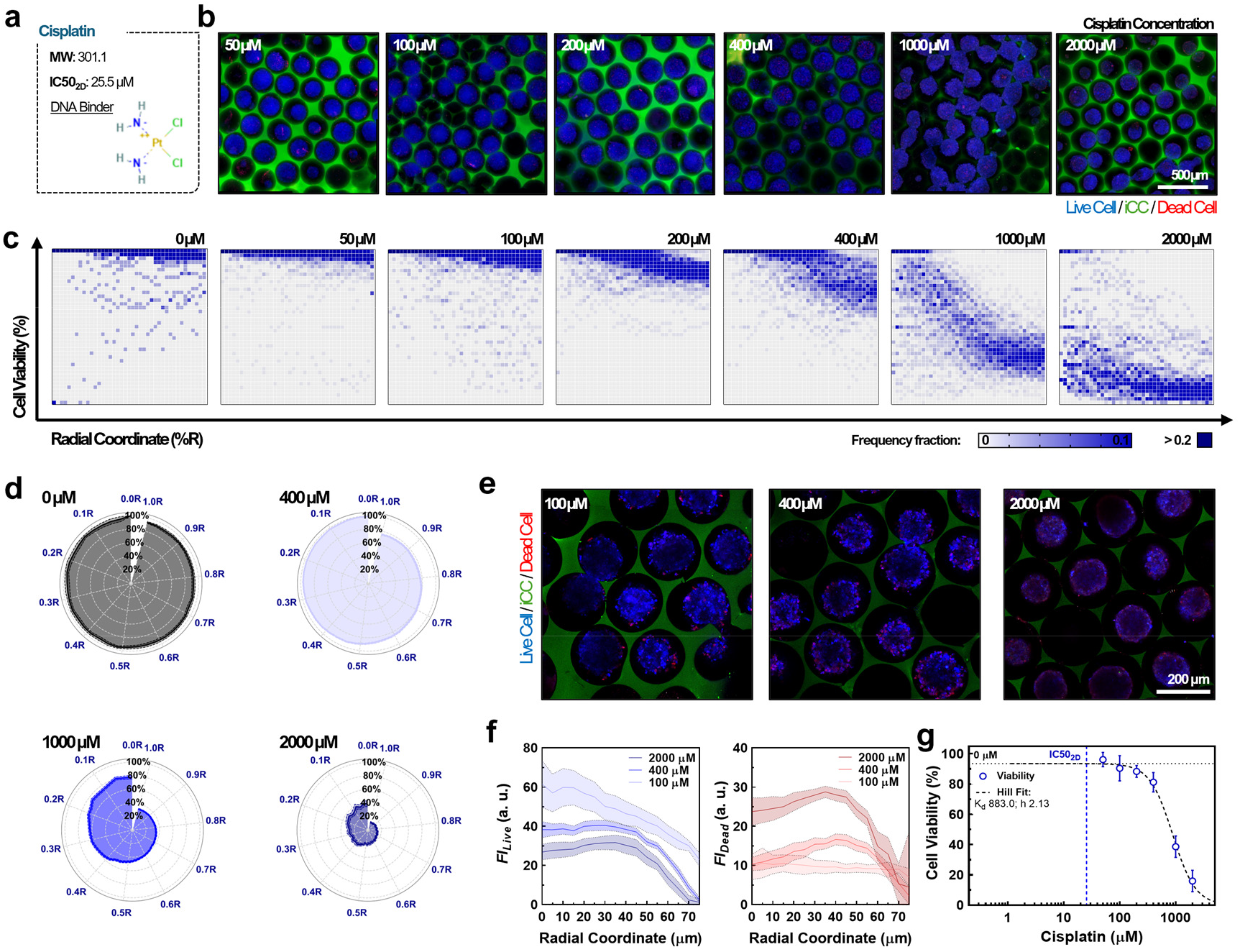
HCHT spatial viability profiling of HepG2 spheroids exposed to cisplatin in the iCC spheroid array (24 h treatment following 6-day spheroid culture). **(a)** Descriptive summary of cisplatin. **(b)** Representative TP-CLSM images showing live (blue) and dead (red) HepG2 cell distributions in response to increasing cisplatin concentrations (50–2000 µM). Green: iCC framework. Images represent sum slice-based *Z* projections. **(c)** Heatmaps of cell viability frequency across radial coordinate (%R) and viability percentile at varying doses. **(d)** Polar plots visualizing spatial viability distributions at 0, 400, 1000, and 2000 µM, showing dose- and depth-dependent reductions in live cell populations. **(e)** Single-slice TP-CLSM images showing radial viability changes at 100, 400, and 2000 µM. **(f)** Radial fluorescence intensity profiles for live (left) and dead (right) cell signals, indicating enhanced viability changes at the spheroid edges. **(g)** Dose–response curve with IC50 estimation *via* Hill fitting.

Next, we explored the spatial cytotoxic response of HepG2 spheroids to paclitaxel, a microtubule-stabilizing agent with a relatively large molecular weight (∼854 Da) and poor aqueous solubility (**Figure 5a**). Unlike doxorubicin and cisplatin, paclitaxel is known as microtubule stabilizer which has *fC50*_*2D*_ as 4.06 µM with 24 h treatment on HepG2 in 2D conventional culture.^70^ For the iCC spheroid array treatment, we chose higher concentration ranges of paclitaxel like cisplatin cases, starting from 10 µM up to 1000 µM—the solubility- limited maximum. Following 24 h exposure, sum-sliced TP-CLSM imaging (**Figure 5b**) revealed minimal red signal accumulation even at the highest concentration, indicating limited cell death across the array. Blue live-cell signals remained dominant in all cases, highlighting the distinct spatial efficacy of paclitaxel. Viability frequency heatmaps (**Figure 5c** and **Figure S8**) showed that, unlike doxorubicin or even cisplatin, paclitaxel did not induce sufficient cytotoxicity at any dose at spheroid edge. Instead, the highest response was limited to ∼50% viability reduction at the spheroid edge, with only marginal effect on the spheroid core. Polar plots of spatial viability (**Figure 5d** and **Figure S9a**) demonstrated consistent but modest viability drop (*i*.*e*., *eViability* _*1*.*0R*_ *− eViability* _*0*.*0R*_) across all doses with 40% viability drop in maximum (**Figure S9b**), reinforcing that cell death was largely localized at peripheral regions and barely appeared in the center of the spheroids. This radial viability plateau across the tested dose range suggests either insufficient paclitaxel transport or a mechanistically delayed cytotoxic onset not captured within the 24 h window. Single-slice TP-CLSM images at representative doses (**Figure 5e**) displayed a unique “ringing” effect, where peripheral red signal encircled a large viable core. This phenomenon became more pronounced at medium and high concentrations, yet no dead cell distribution was observed in the core of spheroids. *Ff* profiles across radial coordinates (**Figure 5f**) further supported this, showing decreasing live-cell signal with dose escalation, but the dead-cell signal remained largely unchanged and confined to the spheroid edge. This stands in contrast to *Ff*_*B*_ and *Ff*_*R*_ patterns observed in the doxorubicin and cisplatin cases and suggests a fundamentally different cell response to paclitaxel which should be further explored. Quantitative dose–response analysis (**Figure 5g**) based on live signal volume showed a shallow slope, requiring extrapolation for Hill equation fitting. The extrapolated *fC50*_*3D*_ value (>>1000 µM) was not practically achievable within the solubility or viability limits of the platform. This result further emphasized the advantage of spatial profiling for drug screening in 3D spheroid platforms: while standard bulk viability metrics may underestimate partial peripheral responses, spatial maps revealed localized drug action that would otherwise be missing.

**Figure 5.**
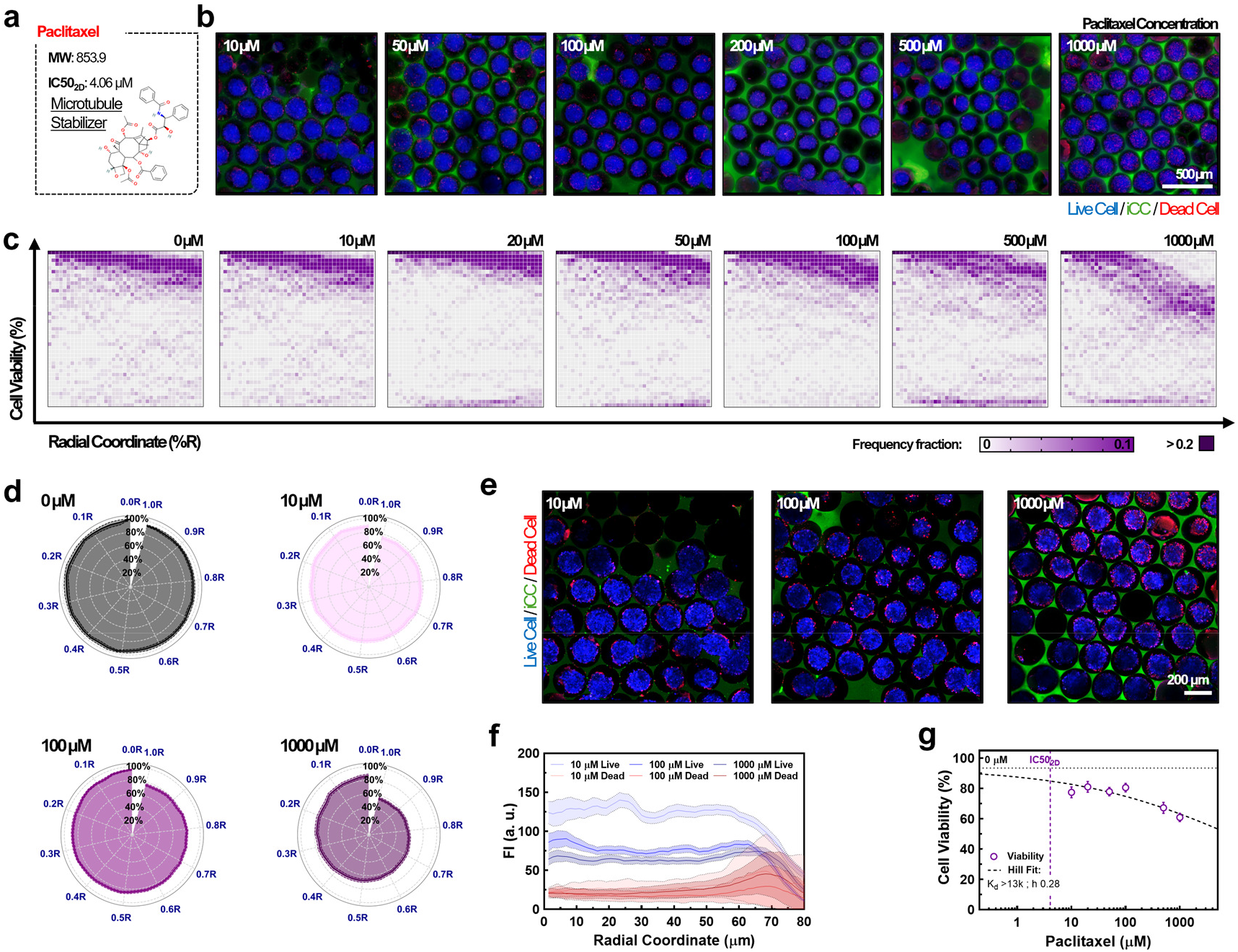
HCHT spatial viability profiling of HepG2 spheroids exposed to paclitaxel in the iCC spheroid array (24 h treatment following 6-day spheroid culture). **(a)** Descriptive summary of paclitaxel. **(a)** Representative TP-CLSM images showing live (blue) and dead (red) HepG2 cell distributions in response to increasing paclitaxel concentrations (10–1000 µM). Green: iCC framework. Images represent sum slice-based Z projections. **(b)** Heatmaps of cell viability frequency across radial coordinate (%R) and viability percentile at varying doses. **(c)** Polar plots visualizing spatial viability distributions at 0, 10, 100, and 1000 µM, showing dose- and depth-dependent reductions in live cell populations. **(f)** Single-slice TP-CLSM images showing radial viability changes at 10, 100, and 1000 µM. **(e)** Radial fluorescence intensity profiles for live (left) and dead (right) cell signals, showing minimal increase in dead cell signal and a significant reduction in live cell signal at higher doses. **(d)** Dose–response curve with IC50 estimation *via* Hill fitting.

## 3. Discussion

In this study, we presented an integrated platform that combined two key components. First, a bioinert iCC hydrogel framework was used to generate high-yield spheroid arrays arranged in a crystalline pattern (*i*.*e*., iCC spheroid array). Second, we developed an automated, time-efficient HCHT image processing pipeline to enable robust spatiotemporal profiling. This platform overcomes the key challenges in 3D culture systems for HCHTS by implementing high-yield spheroid culture with a fully automated data processing pipeline with both physiological relevance and statistical reliability. It addresses challenges like variability in spheroid uniformity and yield,^13^ limited throughput in analysis,^40,41^ manual processing for spheroids identification,^39,42,44^ and limited spatial preciseness and resolution.^39,42,43^ By structurally guiding cellular self-assembly within a precisely defined microarchitecture, the iCC framework enabled uniform spheroid size distribution (*e*.*g*., <10% standard deviation; S.D.) at high yield (*e*.*g*., >50 spheroids·μL^−1^), facilitating reproducible and large-scale data acquisition. The computational pipeline along with the iCC framework significantly enhances throughput and objectivity compared to conventional manual or semi-automated workflows,^15,71^ which are often time-consuming, user-biased, and poorly scalable. Our system enables rapid ROI recognition and extraction, fluorescence-weighted centroiding, multi-parametric signal profiling at the individual spheroid level, and downstream data sorting in spatial coordinate, in <5 sec·image^−1^ with 40—60 spheroids·image^-1^).

The ability of our system to support 3D HCHT spatiotemporal profiling was clearly demonstrated through the analyses of doxorubicin, sEVs, and immune cell dynamics within the iCC spheroid array. For instance, doxorubicin penetration was captured in a dose-dependent manner, transitioning from peripheral restriction at low concentrations to uniform penetration at higher doses. Such dose-dependent penetration profiles align the broader challenges of drug diffusion in solid tumors, where structural barriers like dense extracellular matrices restrict therapeutic access, particularly at sub-saturating concentrations.^72^ In contrast, sEVs exhibited persistent peripheral confinement even after 72 h, indicating their limited transport within dense 3D tumor-like tissues potentially due to their size (∼100 nm). This observation aligns with prior reports on size-dependent limitations of nanoparticle penetration in tumor spheroids,^73^ suggesting that effective nanoparticle-based drug delivery into solid tumors, including sEVs, may require active transport mechanisms. Interestingly, despite the significantly larger size (∼10 µm) of immune cells, our platform revealed distinct cell-type-specific transport behaviors, showing limited penetration of differentiated macrophages (THP-1m) but progressive migration of monocytes (THP-1) throughout the whole spheroid region starting from Day 3. This observation is consistent with the concept of monocyte hijacking,^74^ wherein tumors recruit circulating monocytes and enable their accumulation within the tumor microenvironment.^75,76^ While such infiltration patterns have been previously observed using conventional 3D culture models,^21^ our 3D HCHT platform enables scalable, resolved spatial profiling of immune cell migration dynamics, supporting the potential of our pipeline for investigating tumor–immune interactions and highlight the active migratory potential of monocytes in 3D tumor environments. Together, these findings validate our iCC-based 3D HCHTS platform as a versatile and statistically reliable tool for spatially resolved profiling of various therapeutic agents and cell types in physiologically relevant tumor environments.

To further enhance the practical utility of our platform for comparative drug response analysis, we integrated HCHT viability quantification across multiple chemotherapeutic agents (**Figures 3–5**). Unlike the raw intensity-based spatial tracking used in earlier profiling (**Figure 2**), this phase focused on biologically interpretable endpoints by collecting normalized viability signals from each individual spheroid. As summarized in **Table 1**, spatial viability profiling of HepG2 spheroids exposed to doxorubicin, cisplatin, and paclitaxel revealed critical differences in cytotoxicity profiles within dense 3D tumor-like tissues. Doxorubicin exhibited significant dose-dependent spatial toxicity, especially at higher concentrations, where near-complete cell death occurred across the spheroid volume (**Figure 3**). In contrast, cisplatin, a platinum-based chemotherapeutic, showed higher peripheral cytotoxicity, with lower efficacy in the core region of spheroids even at high doses (**Figure 4**). This behavior is potentially due to its reactivity with a variety of biological molecules (*e*.*g*., nucleic acids, proteins, ECM, and cellular organelles), which slows its diffusion through both the intracellular space and spheroid structure, resulting in a pronounced viability gradient from the core to the periphery. Paclitaxel, with its large molecular weight and poor solubility, induced minimal cell death even at high concentrations, with effects largely confined to the spheroid edge, suggesting limited transport or delayed cytotoxic onset (See **Figure 5**). Interestingly, upon revisiting the raw intensities of live and dead cell signals, we observed a decrease in the live cell signal. However, there was no corresponding increase in the dead cell signal. Notably, the dead cell signal remained confined to the spheroid edge. These observations suggest a distinct mechanism of action for paclitaxel in 3D spheroids, potentially involving hypoxia-induced paclitaxel resistance^77–79^ and hypoxia-induced membrane leakage.^80,81^ These results emphasize the importance of spatial profiling in assessing drug efficacy in 3D systems, where traditional bulk viability measurements may overlook localized effects, particularly at the spheroid periphery. The differences in spatial drug responses further emphasize the need for tailored drug delivery strategies that account for the unique challenges of tissue-like 3D environments, particularly with respect to drug diffusion and localized efficacy.

**Table 1.**
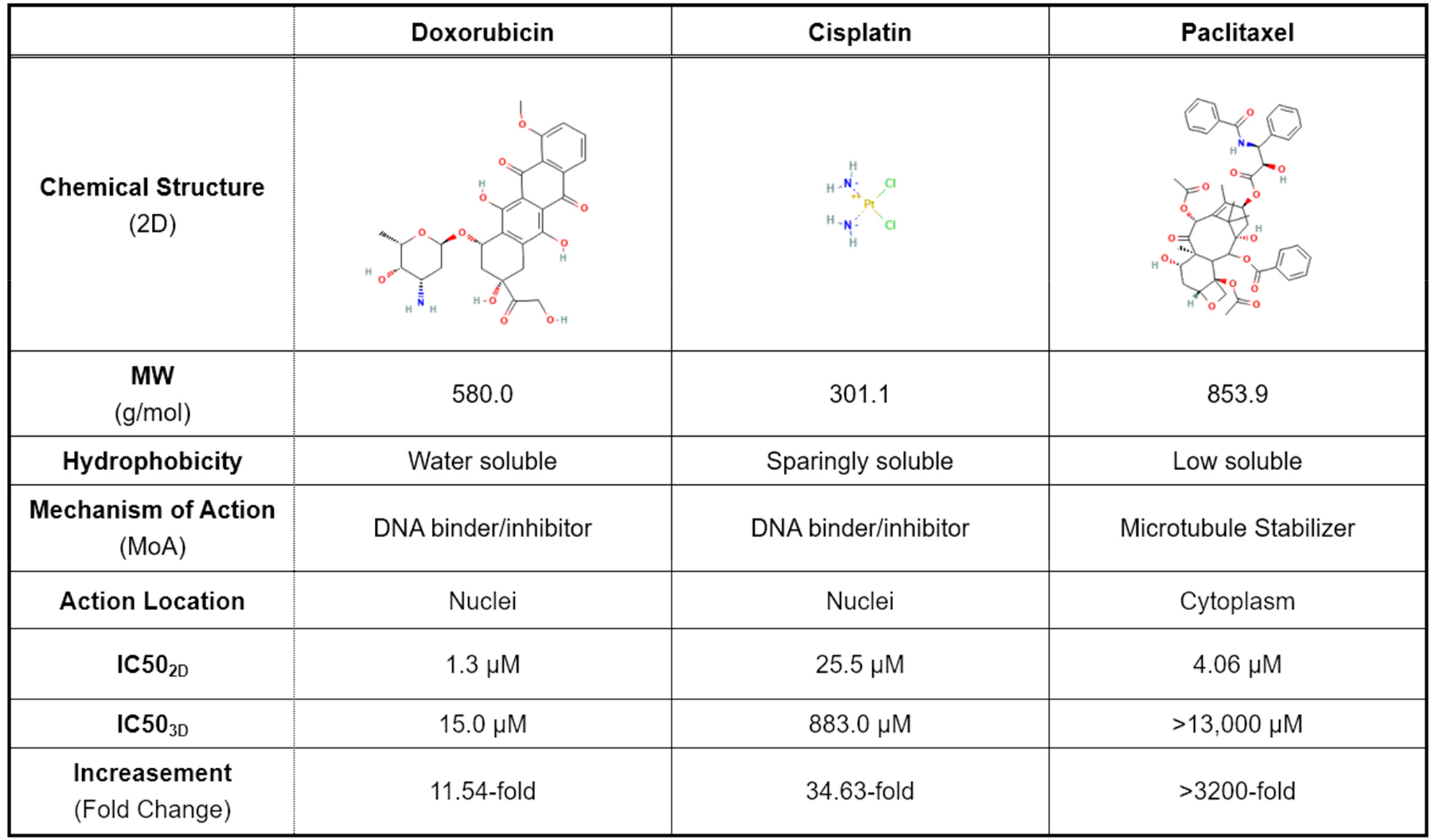
Summary of chemotherapy drug testing with iCC spheroid array.

|  | Doxorubicin | Cisplatin | Paclitaxel |
| --- | --- | --- | --- |
| Chemical Structure (2D) |  |  |  |
| MW (g/mol) | 580.0 | 301.1 | 853.9 |
| Hydrophobicity | Water soluble | Sparingly soluble | Low soluble |
| Mechanism of Action (MoA) | DNA binder/inhibitor | DNA binder/inhibitor | Microtubule Stabilizer |
| Action Location | Nuclei | Nuclei | Cytoplasm |
| IC50<sub>2D</sub> | 1.3 μM | 25.5 μM | 4.06 μM |
| IC50<sub>3D</sub> | 15.0 μM | 883.0 μM | >13,000 μM |
| Increase (Fold Change) | 11.54-fold | 34.63-fold | >3200-fold |

Beyond spatial viability assessment, our platform is expected to offer next-generation HCHT capacity for multiparametric phenotypic readouts across hundreds of individual 3D tumor spheroids in a minute scale, significantly enhanced the processing time from conventional works. This scalability allows robust statistical power and meaningful comparisons between therapeutic agents, concentrations, and treatment durations. The modular nature of our platform is readily adaptable for co-culture models, timelapse studies, and HCHT imaging of additional axes (*e*.*g*., proliferation, apoptosis, hypoxia, or intercellular infiltration), expanding its translational relevance for preclinical drug screening. Furthermore, the standardized ROI extraction abilities and spatial signal outputs generated by our pipeline are inherently compatible with artificial intelligence (AI) and machine learning (ML) modeling as a reliable input dataset. This creates opportunities for predictive modeling of drug efficacy, spheroid phenotype clustering, and automated treatment stratification. In future applications, AI-driven multiparametric integration may enable rapid phenotypic fingerprinting of drug responses, enhancing personalized medicine pipelines and accelerating discovery of effective therapeutic combinations for solid tumors.

## 4. Conclusion

In this study, we developed a versatile 3D HCHT profiling platform, integrating an iCC hydrogel framework with an automated image analysis pipeline. This platform addresses key challenges in 3D HCHT screening systems, including low spheroid uniformity, limited yield, and time-consuming, user-biased manual data processing. By guiding cellular self-assembly within a defined microarchitecture, the iCC framework enables spontaneous formation of crystalline-arranged spheroids (∼79.8 spheroids/mm^2^) with highly uniform size distribution (<10% S.D.), facilitating reproducible, large-scale data acquisition (∼40 spheroids per TP-CLSM image). Complementarily, our automated pipeline enables rapid, unbiased analysis—featuring ROI identification, fluorescence-weighted centroiding, and multi-parametric spatial signal profiling at the individual spheroid level—all completed in under 5 seconds per image with ∼40 spheroids. The utility of this platform was demonstrated through spatial profiling of doxorubicin, evaluation of sEV-based nanocarrier penetration, and characterization of immune cell (THP-1 and THP-1m) dynamics penetrating into spheroids. Our system successfully captured dose-dependent doxorubicin penetration, highlighted sEV transport limitations in 3D tumor spheroids, and revealed distinct immune cell behaviors in 3D tumor spheroids—validating its robustness and potential for profiling and assessment of wide range of therapy in 3D tumor models with high statistical reliability. Moreover, comparative drug response (*i*.*e*., cell viability) profiling uncovered chemodrug-specific cytotoxicity patterns, highlighting the importance of spatial viability assessment in 3D contexts. Scalable and adaptable, the platform integrates high-throughput capacity with automated spatial analysis to support multi-parametric phenotypic profiling across hundreds of spheroids. Its broad applicability—from co-culture models and time-lapse imaging to drug screening and tumor biology studies—suggests its potential as a valuable tool for advancing 3D *in vitro* models in diagnostics, therapy development, and tissue engineering.

## 5. Experimental Section

### 5.1 Materials and Reagents

Alginic acid sodium salt (180947), calcium chloride dihydrate (CaCl^2^; 223506), sodium hydroxide anhydrous (NaOH; S5881), and alginate lyase powder (A1603) were purchased from Sigma-Aldrich (MO). Cryopres dimethyl sulfoxide (DMSO; 092780148) was purchased from MP Biomedicals (CA). Certified^™^ low melt agarose (agarose; 161-3111) was purchased from Bio-Rad (CA). 5-[(4,6-Dichloro-triazin-2-yl)amino]fluorescein hydrochloride (5-DTAF; 21811-74-5) was purchased from Chemodex (Switzerland). Sylgard™ 184 silicone elastomer (polydimethylsiloxane, PDMS; 102092-312) was purchased from VWR (PA).

Minimum essential medium Eagle 1 X (MEM; 10-010-CV) and phosphate-buffered saline 1 X (PBS; 21-040-CM) were purchased from Corning (NY). RPMI-1640 Medium with L-glutamine (RPMI-1640; 112-023-101) was purchased from Quality Biological Inc (MD). Avantor Seradigm USDA-approved origin fetal bovine serum (FBS; 1300–500) was purchased from VWR (PA), kept at temperature of −20 °C. Antibiotic antimycotic (15240096) was purchased from Fisher Scientific (MA). Gibco™ Trypsin-EDTA (25200072) was purchased from Thermofisher Scientific (MA). Phorbol 12-myristate 13-acetate (PMA; P8139) was purchased from Sigma-Aldrich (MO). 4% Paraformaldehyde in 0.1 M phosphate buffer (15735-50S) was purchased from Electron Microscopy Sciences (PA). Cell Explorer^™^ Live Cell Tracking Kit (22620) were purchased from AAT Bioquest (CA). CytoTrace™ Red CMTPX (22015) and Cell Explorer^™^ Live Cell Tracking Kit (22620) were purchased from AAT Bioquest (CA). Vybrant Multicolor Cell-Labeling Kit (DiD; V22889) was purchased from Thermofisher Scientific (MA). BOBO-3 Iodide (570/602) was purchased from Thermofisher Scientific (MA). Doxorubicin Hydrochloride (DOX; D4193) was purchased from Tokyo Chemical Industry Co. (Tokyo, Japan). Cisplatin (89150-634) and Paclitaxel (103607-762) were purchased from VWR (PA).

### 5.2 Bioinert Agarose Hydrogel-Based iCC Framework Fabrication

The iCC hydrogel framework was fabricated based on our previously established method,^48^ with minor modifications to enhance scalability and uniformity for high-throughput applications. Monodisperse alginate microgels were generated via electrospraying of a 2.5%_w/v_ alginic acid solution crosslinked in 200 mM CaCl_2_ using a Spraybase® system. For fluorescence labeling, agarose was conjugated with 5-DTAF under basic conditions. Briefly, hydrated agarose was reacted with 5-DTAF (10 mM in DMSO) and NaOH (1 M aqueous solution), followed by incubation, centrifugation, and multiple water washes. The labeled agarose was mixed at 10%_w/w_ with unlabeled agarose and thermally processed to form a moldable hydrogel. The resulting hydrogel was sliced and reshaped into 1.5 mm-thick disks using temperature cycling within a glass-bottom dish mold. To assemble the microgel template, 6.0 × 10^4^ alginate microgels were suspended in water and deposited into a glass-bottom dish. Gentle sonication removed excess water, enabling formation of an HCP microgel array. The agarose hydrogel was cast over the array, covered with a coverslip, and heated at 70°C for 30 min to ensure full embedding, followed by cooling at room temperature to solidify infiltrated agarose hydrogel. The embedded microgels were removed by sequential enzymatic and ionic degradation, including incubated with alginate lyase (1 mg·mL^−1^) at 37 °C for 24 h, then thoroughly washed with water. Remaining Ca^2^? ions were chelated with repeated PBS (200 mL) washes five times with 1 h interval, occurring the hydrogel turned transparent. The final hydrogel-based iCC framework was rinsed and UV-sterilized for cell seeding and spheroid generation.

### 5.3 Cell Culture, Differentiation, and sEV Collection

Human hepatocellular carcinoma (HepG2; HB-8065, ATCC, VA) cells were cultured in MEM supplemented with 10% fetal bovine serum (FBS) and 1% antibiotic-antimycotic (AA), maintained at 37 °C with 5% CO_2_ in a humidified incubator (MCO-15AC; Sanyo, Japan), and routinely passaged (passages 6–12) to ensure exponential growth. Once HepG2 cells reached >80% confluency, they were washed with PBS, trypsinized, and resuspended at 1.0 × 10_7_ cells·mL^−1^. Human leukemia monocytic (THP-1; TIB-202) [ATCC, VA] cells were cultured in RPMI-1640 supplemented with 10%FBS and 1% AA, maintained at 37 °C with 5% CO_2_ in a humidified incubator (MCO-15AC; Sanyo, Japan), and routinely passaged (passages 6–12) to ensure exponential growth. Following the established THP-1 differentiation protocol,^57^ THP-1 monocytes were differentiated into macrophages by 24 h incubation with 150 nM PMA in T25 flask, followed by 24 h incubation in fresh RPMI medium. For red cell tracker staining of THP-1 lines, 1.0 × 10^6^ mL^-1^ of suspending cells were treated with 5 µM of CMTPX within 2 mL of the cell culture medium for 1 h, followed by washing with PBS and resuspension into 1.0 × 10^6^ mL^-1^ in 1 mL fresh media.

The sEV derived from HepG2 (*i*.*e*., Hep-sEV) was collected following our established method.^61,62^ Once HepG2 cells covered 70−80% of the flask, the cell culture medium was changed into serum-free medium (*e*.*g*., without FBS) after washing three times with PBS. The serumfree medium was collected after 48 h. Then, the medium was filtered with Pore Size 0.22 μm Vacuum Filtration Systems (10040-460; VWR, Radnor, PA) to remove undesired large debris. The medium was filtered through 0.05-μm-pore-size hydrophilic membranes (111103; Cytiva, Marlborough, MA) under mild-vacuum filtration conditions for 2 days. Then, the medium was washed with PBS and concentrated in 100 kDa ultrafiltration centrifugal devices (Spin-X UF Concentrator; Corning, NY). The resulting sEVs were stained with DiD dye by adding 2 μL of DiD to 200 μL of 4.8 × 10^9^ sEVs·mL^-1^ for 10 min in 37°C. The stained sEVs are washed with PBS and concentrated in 100 kDa ultrafiltration centrifugal devices for three times.

With negative staining by UranyLess (22409; Electron Microscopy Sciences, PA), the morphology and spherical nanostructure of sEVs were observed with TEM (JEOL 2011; JEOL Ltd., Tokyo, Japan). Nanoparticle Tracking Analysis (NTA; NanoSight LM10 system; Malvern Instrument Ltd., U.K.) was performed using diluted sEVs (100 times dilution; 10 μL in 1 mL of PBS buffer) and analyzed with the NTA 3.3 analytical software suite. The surface charge of the sEVs was measured using a Zetasizer (Malvern Zetasizer Nano ZS; Malvern Instruments Ltd., U.K.). For each sample, five measurements were performed, with 30–100 autoruns per measurement, using the Hückel model. sEV marker expressions of CD9, CD63, and CD81 (EXOAB-KIT-1; System Biosciences, CA) were identified through Western blot analysis under a bioimaging system (C400 Bioanalytical Imager; Azure Biosystems, CA).

### 5.4 Spheroid Array Generation within the iCC Framework

For live tracking, cells were incubated with 20 µM Cell Explorer™ Live Cell Tracking dye in 2 mL of culture medium for 30 min, then washed and resuspended in fresh medium at the same concentration. To seed the iCC framework, 4.5 mL of fresh medium was added to the framework positioned inside a perpendicular-designed, PDMS-molded 50 mL centrifuge tube. A 250 µL cell suspension containing 1.5 × 10_7_ cells was gently mixed with the surrounding medium, followed by centrifugation at 400 rpm for 3 min. The resulting cell pellet outside the framework was gently redistributed around the structure. This process was repeated to achieve a total of 3.0 × 10_7_ cells per framework. The seeded framework was then transferred to a non-treated 60 mm culture dish containing 5 mL of fresh medium and incubated for 6 days at 37 °C with 5% CO2, with medium changes every 48 h. All drug and transport assessments were conducted post day 6.

### 5.5 Data Acquisition for Small Molecule, sEV, and Cell Transport Evaluation

To assess molecular transport, doxorubicin was dissolved in DMSO (10 mM) and diluted into HepG2 medium at 0 (control), 10, 50, and 200 μM. iCC spheroid arrays were incubated with doxorubicin-containing media for 24 h. BOBO-3 Iodide was then applied in fresh medium for 15 min to stain dead cells. For sEV penetration, 2 mL of DiD-labeled sEV-containing media (2.4 × 108 sEVs·mL^−1^) was added and incubated for 24, 48, and 72 h. To evaluate THP-1 cell infiltration, 2 mL of CMTPX-stained THP-1 suspensions (1.0 × 10^6^ cells·mL^−1^), both undifferentiated and PMA-differentiated (macrophage-like), were added and incubated for 3, 5, and 7 days. After each treatment, arrays were fixed with 2% paraformaldehyde (PFA) for 3 h, washed with PBS, and stored in 2 mL PBS. Multiphoton imaging was performed using the Leica Stellaris 8 DIVE Point Scanning Confocal Microscope across three fluorescence channels: blue (live HepG2), green (doxorubicin or 5-DTAF in iCC framework), and red (dead cells, sEVs, or THP-1 cells).

### 5.6 Data Acquisition for Chemotherapy Drug Treatment and Assessment

To evaluate chemotherapeutic efficacy, doxorubicin was dissolved in DMSO (20 mM) and diluted to 0 (control), 5, 20, 50, 100, 200, and 500 μM. Cisplatin (10 mg·mL^−1^, ∼33.2 mM in DMSO) was diluted to 50, 100, 200, 400, 1000, and 2000 μM. Paclitaxel (50 mg·mL^−1^, ∼58.6 mM in DMSO) was diluted to 10, 50, 100, 200, 500, and 1000 μM. iCC spheroid arrays were incubated with each drug for 24 h, followed by BOBO-3 Iodide staining for 15 min. Multiphoton imaging was performed using the same confocal setup, capturing blue (live cells), green (5-DTAF in framework), and red (dead cells) channels.

### 5.7 Automated Data Processing Algorithm Workflow and Employment

To enable high-throughput, quantitative spatial analysis of 3D spheroid arrays, we developed an all-in-one automated data processing pipeline in Python. This workflow integrates image preprocessing, region-of-interest (ROI) extraction, centroid localization, and spatial signal quantification. Multiphoton z-stack images of spheroid-laden iCC frameworks were acquired using a Leica Stellaris 8 DIVE Point Scanning Confocal Microscope. The datasets comprised three fluorescence channels: blue for live HepG2 cells within spheroids, green for either iCC geometry visualization (*via* 5-DTAF) or oversaturated doxorubicin, and red for markers of dead cells, small extracellular vesicles (sEVs), or THP-1 infiltration. Each channel was separated and converted into grayscale stacks for independent analysis. To isolate individual void regions corresponding to spheroids, a vision-based ROI detection algorithm was applied to the green channel using OpenCV’s Hough Circle Detection method.^53^ The identified circular ROIs were uniformly applied across all channels, enabling channel-aligned, ROI-extracted image stacks for each spheroid. These reconstructed stacks preserved spatial fidelity across fluorescence channels.

For each ROI, spheroid boundaries were delineated by applying intensity thresholding to both the red (*Ff*_*R*_) and blue (*Ff*_*B*_) fluorescence channels, producing thresholded outputs *Ff*_*R,trs*_ and *Ff*_*B,trs*_. These were then combined using a logical OR operation to generate a composite spheroid mask *Ff*_*RUB,trs*_(*X*_*i*_, *y*_*j*_, *Z*_*k*_), enabling contour identification independent of cell viability. The spheroid centroid 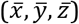 was determined in two steps. First, the *Z*-coordinate was identified as the image slice (Z-plane) with the largest projected spheroid area, representing the maximal cross-sectional signal and minimizing potential bias from light penetration artifacts (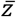 **Equation 1**). Subsequently, the *X*- and *Y*-coordinates were computed based on the fluorescence intensity-weighted average of the thresholded composite mask (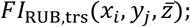 **Equation 2**), enabling accurate localization of the spheroid center in 3D space.

Using the identified centroids, pixel-wise radial distances were vectorized and computed, and signal intensities from each channel were collected radially outward (**Equation 3**). Two modes of quantification were implemented: (i) absolute intensity mapping and (ii) thresholded intensity mapping for categorical analysis (*e*.*g*., dead cell presence-mediated cell viability; **Equation 4-5**). Extracted spatial data from individual spheroids were sorted, reorganized, and exported as .CSV files for statistical and graphical analysis. Intensity or cell viability distributions were plotted as functions of radial distance to characterize signal penetration and localization. Outputs included spatial signal distribution curves, penetration depth quantification, and cellular viability/toxicity profiles. Detailed descriptions and instructions are available on GitHub (https://github.com/HyunsuJeon-ND/iCC_Automated_HCHT_Radial_Analysis).

## Supporting information

bhiccdrugtesting_SI

## Associated Content

### Supporting Information

Supporting Information is available from the author (“[SI] BhiCC Drug Testing_ACS-AMI_Final.pdf”).

## Author Information

## Author Contributions

H.J. contributed to conceptualization, framework fabrication, spheroid culture, drug testing, computational algorithm development, data analysis, and manuscript drafting. G.K. contributed to small extracellular vesicle isolation, purification, characterization, and staining. J.C. and Y.C. contributed to computational algorithm conceptualization, development, and validation. Y.W. contributed to conceptualization, experimental design, and manuscript editing. All authors discussed the results, provided input, and approved the final manuscript for submission.

## Acknowledgement

We acknowledge funding from NSF Career Award (NSF CBET-2337387). This research was funded in part by a Leiva Graduate Fellowship in Precision Medicine from the Berthiaume Institute for Precision Health at the University of Notre Dame. Hyunsu Jeon received partial support from the Notre Dame Scientific Artificial Intelligence (SAI) Initiative for this work. All the TP-CLSM imaging was carried out in part in the Notre Dame Integrated Imaging Facility, University of Notre Dame, using Leica Stellaris 8 DIVE Point Scanning Confocal Microscope.

