## Supplementary material for "Automated High-Content, High-Throughput Spatial Analysis Pipeline for Drug Screening in 3D Tumor Spheroid Inverted Colloidal Crystal Arrays": bhiccdrugtesting_SI

**This PDF file includes:**

- 9 Supplementary Figures
- 1 Supplementary Table

**Table of Contents**

**Supplementary Figures ············································································ S-2**

**└ Figure S1.** Hepatocarcinoma (HepG2) spheroid size distribution within inverted colloidal crystal (iCC) spheroid array from iCC framework approach. ······································· S-2

**└ Figure S2.** iCC void recognition and region of interest (ROI) designation using Hough Circle Detection (OpenCV). Representative output (left), where identified circular voids are marked as ROIs. The corresponding table (right) lists the pixel coordinates (X–Y) and radii of each detected circle used for downstream analysis.···································································· S-3

**└ Figure S3.** Characterization of HepG2-derived small extracellular vesicles (sEVs). **(a)** Representative transmission electron microscopy (TEM) image. **(b)** Nanoparticle tracking analysis (NTA) showing size distribution. **(c)** Zeta potential measurement indicating surface charge. **(d)** Western blot analysis confirming expression of sEV marker proteins. ···························· S-4

**└ Figure S4.** Individual plots of viability output from doxorubicin-treated iCC spheroid array. ················································································································ S-5

**└ Figure S5.** Polar plots and statistical bar graphs of spatial viability from doxorubicin-treated iCC spheroid array. ························································································ S-6

**└ Figure S6.** Individual plots of viability output from cisplatin-treated iCC spheroid array.··········································································································S-7

**└ Figure S7.** Polar plots and statistical bar graphs of spatial viability from cisplatin-treated iCC spheroid array. ····························································································· S-8

**└ Figure S8.** Individual plots of viability output from paclitaxel-treated iCC spheroid array. ················································································································ S-9

**└ Figure S9.** Polar plots and statistical bar graphs of spatial viability from paclitaxel-treated iCC spheroid array. ···························································································· S-10

**Supplementary Tables ············································································ S-11**

**└ Table S1.** Summary of processing time for each module in the automated analysis pipeline for the iCC spheroid array developed in this study. ····················································· S-11

**Supplementary Figures**

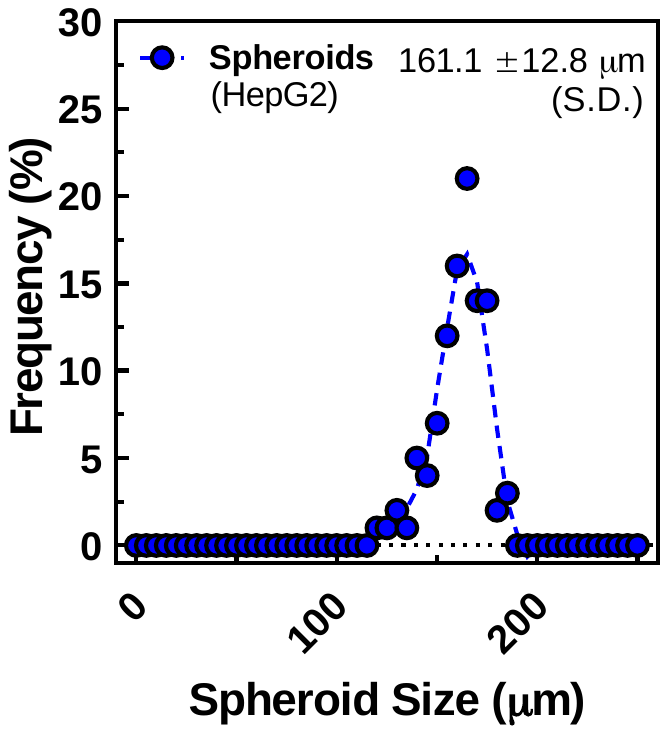

**Figure S1.** Hepatocarcinoma (HepG2) spheroid size distribution within inverted colloidal crystal (iCC) spheroid array from iCC framework approach.

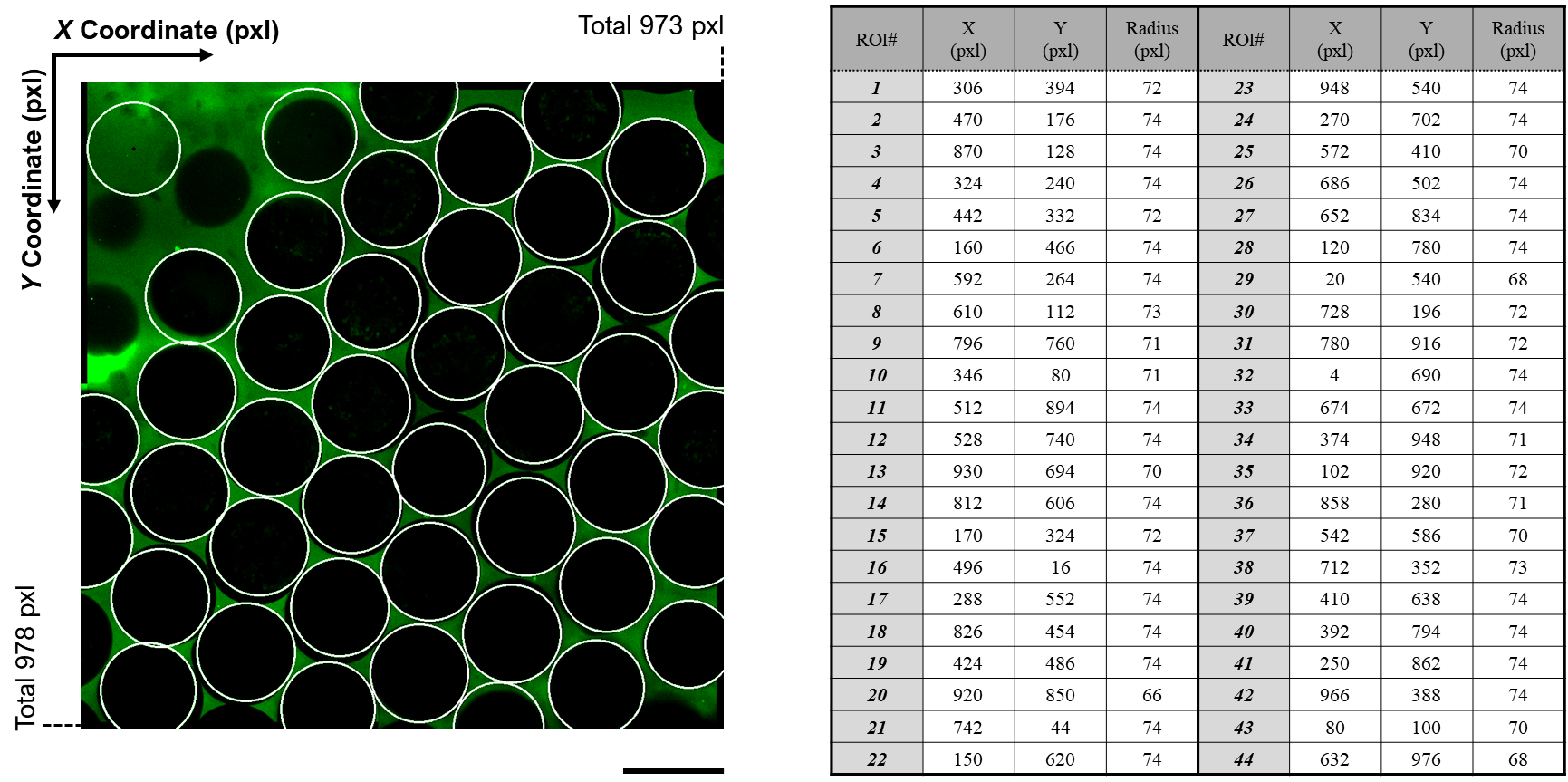

**Figure S2.** iCC void recognition and region of interest (ROI) designation using Hough Circle Detection (OpenCV). Representative output (left; scale bar: 200 µm), where identified circular voids are marked as ROIs. The corresponding table (right) lists the pixel coordinates (X–Y) and radii of each detected circle used for downstream analysis.

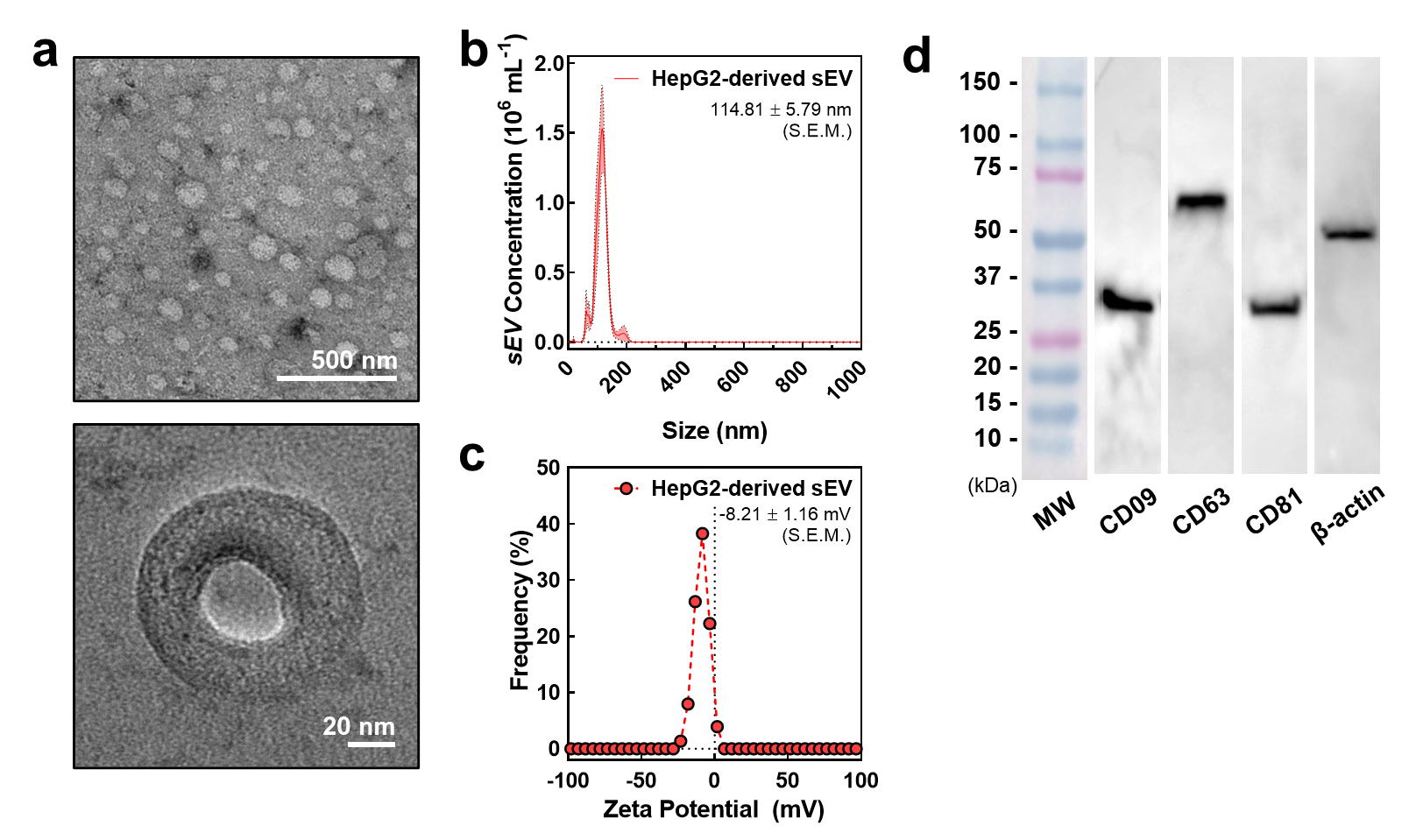

**Figure S3.** Characterization of HepG2-derived small extracellular vesicles (sEVs). **(a)** Representative transmission electron microscopy (TEM) image. **(b)** Nanoparticle tracking analysis (NTA) showing size distribution. **(c)** Zeta potential measurement indicating surface charge. **(d)** Western blot analysis confirming expression of sEV marker proteins.

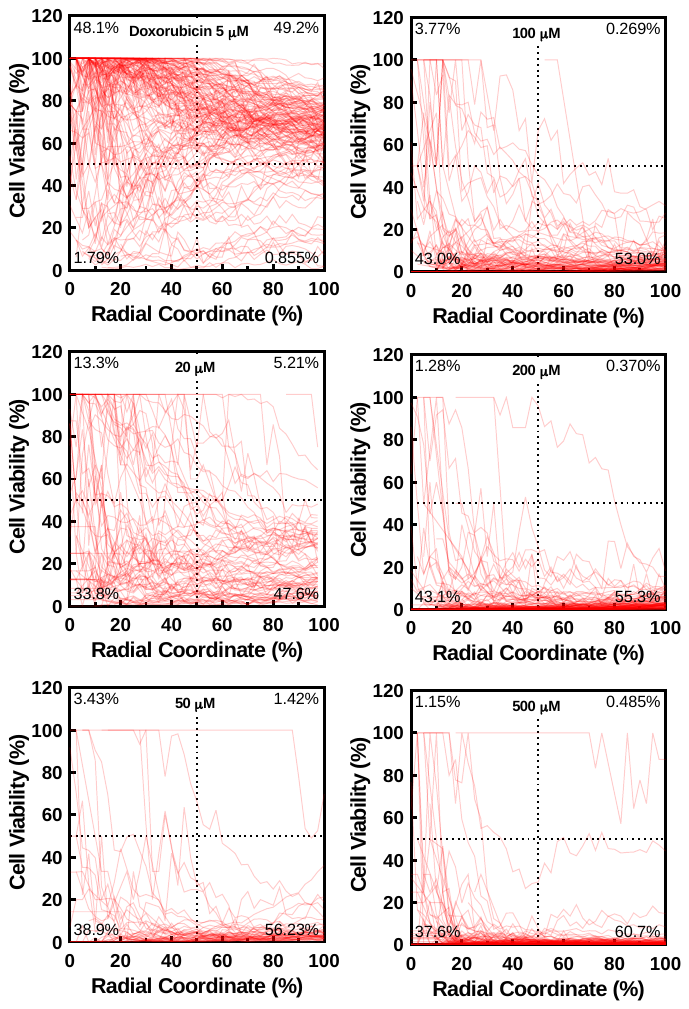

**Figure S4.** Individual plots of viability output from doxorubicin-treated iCC spheroid array.

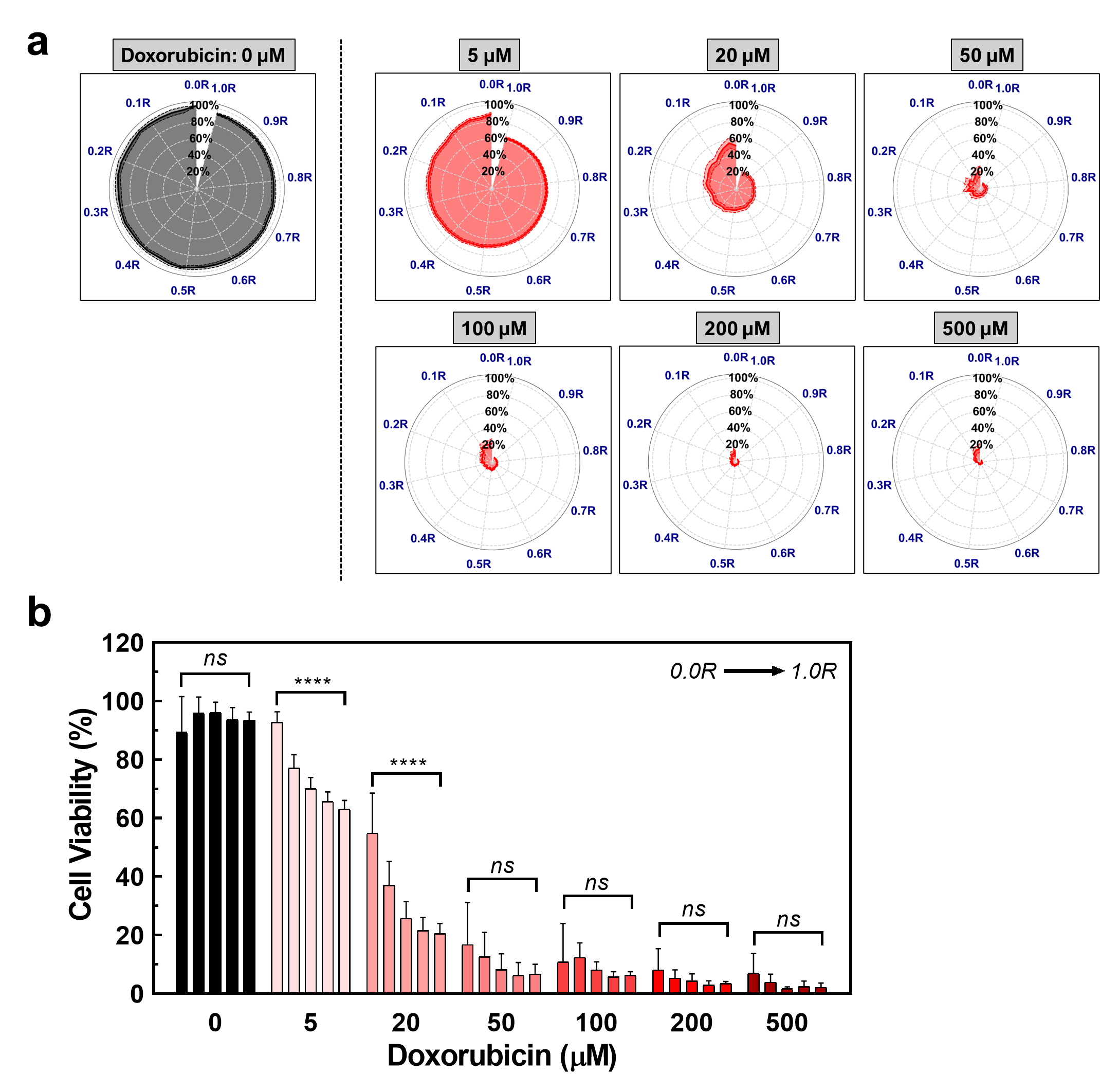

**Figure S5. (a)** Polar plots of spatial viability from doxorubicin-treated iCC spheroid array. **(b)** Spatial viability profiles corresponding to each doxorubicin concentration (0.0R, 0.25R, 0.5R, 0.75R, and 1.0R, respectively). Statistical analysis was conducted using one-way ANOVA across spatial positions (ns: not significant; ****: *P* < 0.0001).

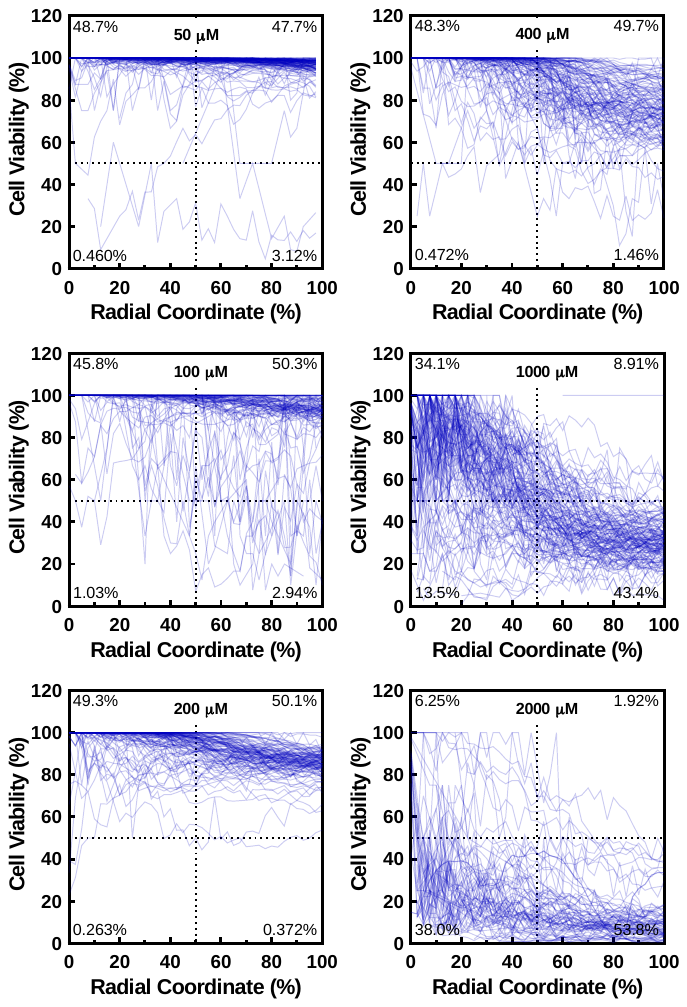

**Figure S6.** Individual plots of viability output from cisplatin-treated iCC spheroid array.

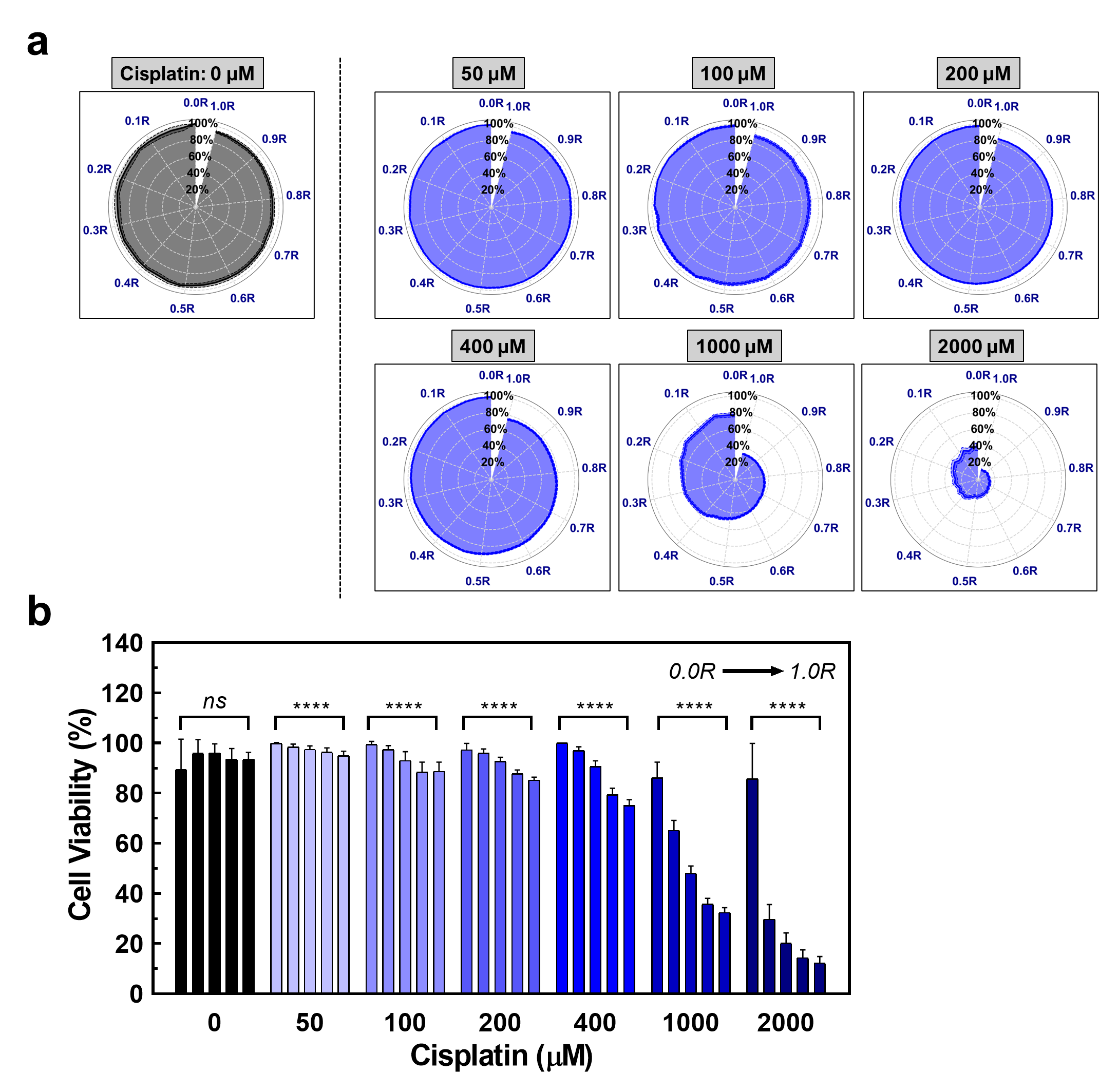

**Figure S7. (a)** Polar plots of spatial viability from cisplatin-treated iCC spheroid array. **(b)** Spatial viability profiles corresponding to each cisplatin concentration (0.0R, 0.25R, 0.5R, 0.75R, and 1.0R, respectively). Statistical analysis was conducted using one-way ANOVA across spatial positions (ns: not significant; ****: *P* < 0.0001).

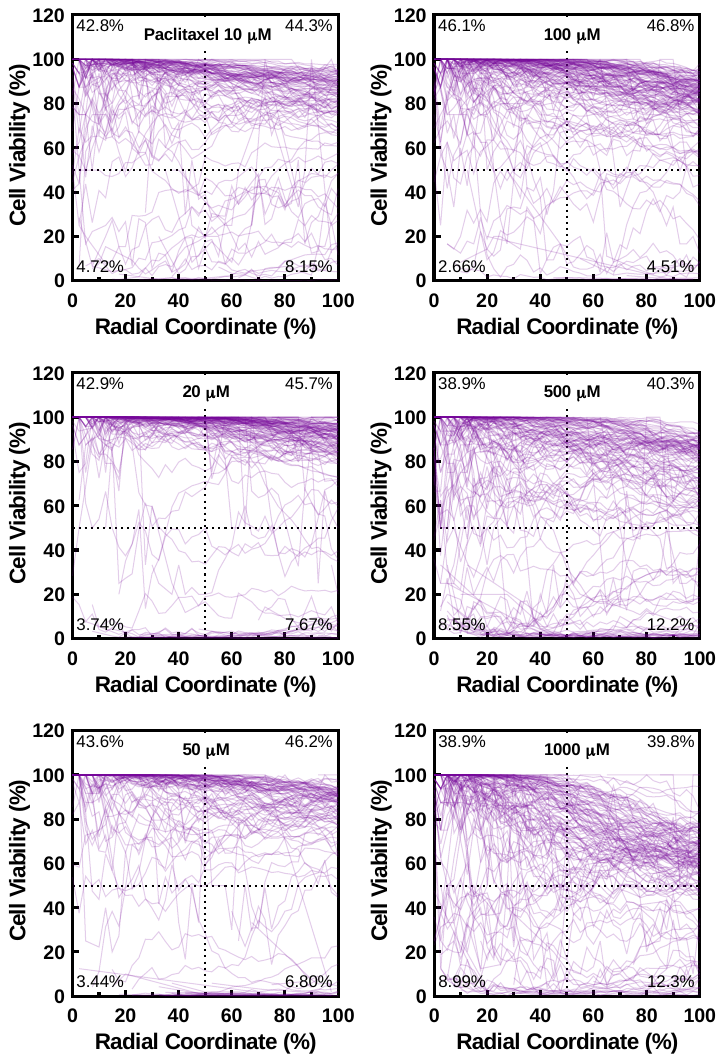

**Figure S8.** Individual plots of viability output from paclitaxel-treated iCC spheroid array.

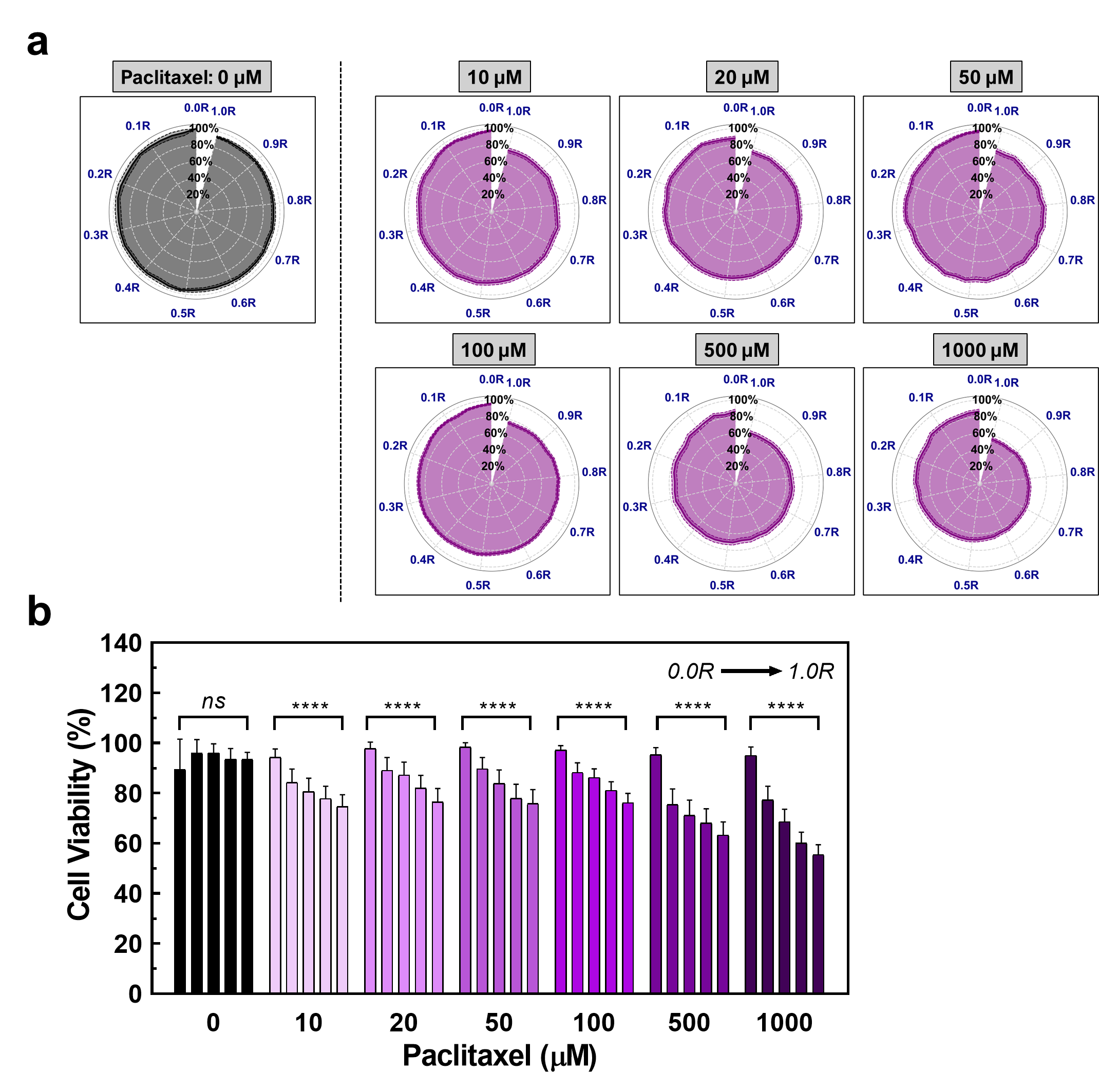

**Figure S9. (a)** Polar plots of spatial viability from paclitaxel-treated iCC spheroid array. **(b)** Spatial viability profiles corresponding to each paclitaxel concentration (0.0R, 0.25R, 0.5R, 0.75R, and 1.0R, respectively). Statistical analysis was conducted using one-way ANOVA across spatial positions (ns: not significant; ****: *P* < 0.0001).

**Supplementary Tables**

Table S1. Summary of processing time for each module in the automated analysis pipeline for the iCC spheroid array developed in this study (*Presented for comparison only; not included in final processing time total).

| **Processing Device** | | **Personal Laptop** | | **Computational Facility (CPU)** | | **Computational Facility (CPU+GPU)** | |
| --- | --- | --- | --- | --- | --- | --- | --- |
| Device Specifications | Central Processing Unit (CPU) | Intel® Core i7-1165G7 | | AMD EPYC 7543 | | Intel(R) Xeon(R) Gold 6326 | |
|  | Core | 4 | | 32 | | 32 | |
|  | Memory (RAM) | 16 GiB | | 256 GiB | | 256 GiB | |
|  | Storage Type | 1TB NVMe SSD | | Networked AFS | | Networked AFS | |
|  | Graphics Processing Unit (GPU) | IntelR Iris Xe Graphics | | Matrox G200eW3 Graphics | | 4 x NVIDIA A10 | |
| Input Image Details | Dimension (pxl) | 960–970, 960–970, 20–30 (*X*, *Y*, *Z*, respectively) | | | | | |
|  | Conversion Vector (µm/pxl) | [*X*, *Y*, *Z*]=[1.354, 1.354, 10] (µm/pxl) | | | | | |
|  | Image Type (TIFF) | 8-bit, 3 channels, 50-80 MB | | | | | |
| **Processing Time (s)** | | **Average** | **S.D.** | **Average** | **S.D.** | **Average** | **S.D.** |
| Region of Interest (ROI) | Recognition | 0.563 | 0.089 | 2.727 | 0.525 | 1.124 | 0.583 |
|  | Extraction | 0.661 | 0.069 | 1.484 | 0.294 | 0.945 | 0.331 |
| Cell Spheroid | Centroid Determination | 0.601 | 0.098 | 0.188 | 0.017 | 0.228 | 0.051 |
|  | *FI Sorting/ Collection  (For-loop-based sorting) | *42.755 | *5.028 | *34.663 | *7.936 | *17.508 | *4.023 |
|  | FI Sorting/ Collection  (Vectorized sorting) | 4.413 | 0.709 | 1.464 | 0.270 | 1.983 | 0.659 |
|  | Post Processing | 0.175 | 0.028 | 0.131 | 0.024 | 0.120 | 0.052 |
| Processing Time Total | | 6.413 | 0.993 | 5.994 | 1.130 | 4.400 | 1.676 |
| ROI Throughput (N image-1) | | 36.250 | 3.832 | 36.250 | 3.832 | 36.250 | 3.832 |
